# Gut colonization by *Bacteroides fragilis* at juvenile age alters microbiota composition and accelerates type 1 diabetes progression in non-obese diabetic mice

**DOI:** 10.1101/2025.06.02.657398

**Authors:** Radhika R. Gudi, Harrison Taylor, Benjamin M. Johnson, Ruchika Maurya, Mary E. Mulligan, Loni Carter, Caroline Westwater, Chenthamarakshan Vasu

## Abstract

Type 1 diabetes (T1D) in humans is associated with higher Bacteroidetes : Firmicutes ratio and higher abundance of Bacteroides genus members. *Bacteroides fragilis* (BF) is an integral component of the human colonic commensal microbiota. Here, we show that gut colonization of specific pathogen-free (SPF) non-obese diabetic (NOD) mice by BF at a juvenile age induces a pro-inflammatory immune response and accelerated disease progression. NOD mice born to BF-monocolonized parents not only showed rapid disease progression compared to germ-free (GF) controls but also preserved accelerated disease onset and higher disease incidence upon conventionalization, suggesting that BF contributes to a pro-inflammatory response and autoimmunity in T1D. Interestingly, we found that while gut microbiota composition was different in control and BF-colonized SPF mice, presence of BF alone could significantly impact the acquisition of microbial communities upon conventionalization of gnotobiotic mice. Bulk RNAseq analysis of colon tissues revealed profound differences in the gene expression pattern of GF and BF-monocolonized mice as well as their conventionalized counterparts, shedding light on the probable mechanisms contributing to accelerated disease onset in mice that are exposed to BF. We found that mucin production is downregulated and the abundance of mucin degraders such as *Akkermansia muciniphila* is profoundly lower in BF-colonized mice that are conventionalized. Overall, these studies demonstrate that early life acquisition of BF-like distal gut commensals could have profound modulatory effects on the eventual overall gut microbiota structure, immune function, and β-cell specific autoimmune outcomes under genetic susceptibility.

## Introduction

While gut commensal microbes contribute profoundly to human health and immune system maturation(1–4), reports also support the notion that gut commensals can contribute to the pathogenesis of many autoimmune diseases, including type 1 diabetes (T1D)(5–11). Initial acquisition of gut microbial communities early on in life can have profound long-term physiological consequences, including susceptibility to autoimmune diseases in at-risk subjects. Studies have shown that the microbiota composition of early life has a role in promoting susceptibility to or protection from diseases such as T1D and metabolic disorders later in life (12–15). Importantly, higher abundances of Bacteroidetes phylum members including *Bacteroides s*pp. have been detected in the gut of not only animal models of T1D, but also in T1D patients and at-risk children who have progressed towards developing disease symptoms(6; 12; 16-22), prompting the suggestion that *Bacteroides* members could exert pro-autoimmune effects under T1D susceptibility. Insufficient exposure to *Escherichia coli* LPS for early life immune system training and maturation in association with overrepresentation of, and exposure to, *Bacteroides* members such as *B. dorei* and *B. fragilis* have been implicated in T1D disease onset in children(13; 22). However, how these gut symbionts contribute to disease progression in T1D is largely unknown.

*Bacteroides fragilis* (BF) is a Gram-negative anaerobe and an integral component of the colonic commensal microbiota in many mammals(23). Although BF represents a very small portion of the gut microbes(24), it is the most commonly isolated bacterium from clinical cases of intra-abdominal abscesses(25; 26), perhaps due to its ability to colonize the colonic crypt(27). While the capsular polysaccharide of BF is necessary for its growth, as well as abscess formation, in the host(26; 28; 29), these components, PSA in particular, have been demonstrated to have the ability to promote immune regulation in a TLR2-dependent manner(27; 30-32). Importantly, BF has been widely studied as a model organism to determine the influence of commensal bacteria in immune regulation(27; 33; 34). Studies using pre-clinical models of various inflammatory diseases have demonstrated the health benefits of this bacterium and its PSA(27; 35-39). The ability of BF to inhibit gut colonization by pathogens has also been demonstrated(40; 41).

In a previous report(42) using heat-killed (HK) wild-type (WT) and polysaccharide A (PSA) deficient BF, we showed that exposure of intestinal and systemic compartments to this bacterium can produce opposing PSA-toll-like receptor 2 (TLR2) interaction dependent effects on autoimmune progression and T1D incidence in the non-obese diabetic (NOD) mouse model. These effects of BF suggested that exposure of the systemic compartment to BF-like gut commensal organisms with immune regulatory properties can result in accelerated autoimmune progression in T1D.

In this report, using T1D-prone NOD mice housed in the conventional/specific pathogen-free (SPF) and germ-free (GF) facilities, we have studied the impact of juvenile age gut colonization by BF on T1D disease incidence, immune function, and overall gut microbiota composition. Early life gut colonization of NOD mice by BF caused the rapid onset of hyperglycemia in both conventional and GF mice. Gut colonization by BF also impacted the host intestinal gene expression profile as well as acquisition of gut microbiota communities upon conventionalization, with notable inhibition of the mucin synthesis pathway in the host as well as acquisition of the mucin-degrading *Akkermansia muciniphila* in the gut microbiota. Overall, these studies suggest that early life acquisition of BF-like distal gut commensals, in excess, could have profound modulatory effects on the overall gut microbiota structure and function, host physiology, and β-cell specific autoimmune outcomes under genetic susceptibility.

## Materials and Methods

### Mice and Bacteria

Wild-type NOD/ShiLtJ (NOD-WT) and NOD-*Scid* mice were purchased from the Jackson laboratory (Maine, USA). Germ free (GF) - NOD mice and NOD-TLR2 knockout (NOD-TLR2 KO) mice were originally provided by Dr. Chervonsky (University of Chicago). Breeding colonies of these strains were established and maintained in the conventional (specific pathogen free; SPF) facility or the gnotobiotic core (germ-free; GF) facility of the Medical University of South Carolina (MUSC).The GF status of gnotobiotic mice were tested routinely by the core facility by culture methods. All animal studies were approved by the Institutional Animal Care & Use Committee (IACUC) of MUSC. To detect hyperglycemia in NOD-WT, NOD-TLR2 KO and NOD-*Scid* mice, glucose levels in blood collected from the tail vein were determined at weekly intervals using the Ascensia Micro-fill blood glucose test strips and an Ascensia Contour blood glucose meter (Bayer, USA). Tail vein blood samples were collected in sodium citrate vials, from NOD mice that were housed in the gnotobiotic isolators bi-weekly and tested for glucose levels using the test strips outside the isolator. In some cases, NOD mice from the gnotobiotic isolators were pulled when they appeared unthrifty with ruffled fur and tested for blood glucose levels to confirm hyperglycemia. Blood glucose level of >250 mg/dl was considered diabetic. In some experiments, mice in the conventional facility were given a cocktail of broad spectrum antibiotics (Abx) as described in our recent report(43) to deplete the gut microbiota. Depletion of gut microbiota was confirmed by culturing fecal pellet suspension on brain heart infusion agar (BHI) plates under aerobic and anaerobic conditions as described before(43).

BF (ATCC 25285; NCTC 9343) was cultured from single colonies in complete BHI medium under anaerobic condition for up to 48h; this initial culture was diluted 50-fold using complete medium and cultured for an additional 16h. Different dilutions were plated on BHI Agar plates and cultured for 24h for determining the colony counts. SPF -NOD mice were given approximately 1x10^10^ CFU (for juvenile) or 1x10^11^ CFU (for adult mice) BF in BHI broth or BHI broth alone (control) for 3 consecutive days by oral gavage. GF-NOD mice were monocolonized by giving 24h cultures from a single PCR confirmed colony by oral gavage. Gut colonization of the NOD parents and litters of the gnotobiotic facility by BF was confirmed by anaerobic cultures of fecal suspension on BHI agar plates followed by PCR and PCR of fecal suspension using BF specific- and universal-16S rRNA gene targeted PCRs. Gut colonization efficiency of SPF mice by BF was assessed by qPCR of fecal DNA preparation for the levels of BF specific- and total-16S rRNA genes.

### Conventionalization of GF and BF-monocolonized mice

About four-week-old female GF-NOD mice and BF-monocolonized NOD mice were pulled from the gnotobiotic isolators, housed on pooled and equally distributed dirty bedding from the breeding cages of conventional facility NOD mice. The bedding material was replaced thrice with fresh dirty bedding from the same breeding cages every other day. To confirm conventionalization, fresh fecal pellets collected from individual mice after 30 days were tested for bacterial colony counts after culturing on BHI agar plates under aerobic and anaerobic conditions. Fresh fecal pellets collected at a 30-day timepoint were used for preparing DNA to determine the abundance of total bacterial- and BF-specific DNA, and microbial community profiling. Cohorts of conventionalized mice were monitored for up to 30 weeks of age for T1D incidence or euthanized at 12 weeks of age to collect pancreatic tissues for determining the degree of insulitis.

### Reagents and in vitro / ex vivo assays

Immunodominant β-cell antigen peptides [viz., 1. Insulin B(9–23), 2. GAD65(206–220), 3. GAD65(524–543), 4. IA-2beta(755–777), 5. IGRP(123–145)] were pooled at an equal molar ratio and used as a β-cell-Ag peptide cocktail as described in our earlier study(44). PMA, ionomycin, Brefeldin A, magnetic bead based multiplexed cytokine assay kits and unlabeled and fluorochrome labeled antibodies, and other key reagents were purchased from BD Biosciences, eBioscience/Thermo/Invitrogen, Millipore-Sigma, Miltenyi Biotec, StemCell Technologies, R&D Systems, and Biolegend.

In some experiments, mice were euthanized at different time points, and tissues (spleen, pancreas, pancreatic LN and distal colon) were collected for various assays. Pancreatic tissues were fixed in 10% formaldehyde; 5-µm paraffin sections were made and stained with hematoxylin and eosin (H&E). Stained sections were analyzed using a 0-4 grading system as described in our earlier report(45). Approximately 50 islet areas/mouse were examined. Spleen and PnLN cells were cultured in the presence of β-cell antigen peptide or anti-CD3 antibody for 24h, and the cell free supernatants were tested for cytokine levels. Luminex technology based multiplex assays were read using FlexMap3D instrument. In some assays, spleen and PnLN cells were subjected to staining for flow cytometry before or after stimulation using PMA/Ionomycin for 4h in the presence of Brefeldin A as described before(42). Flow cytometry data were acquired using the FACS Verse instrument and analyzed by using FlowJo application.

### Quantitative PCR (qPCR) assay

qPCR assay was carried out to determine the expression levels of various proteins as well as fecal bacterial DNA levels as described in our previous reports(42; 43; 46). Briefly, for determining the expression levels of mRNA for various proteins, total RNA preparations were made from the distal colon tissues by employing Trizol method, subjected to cDNA synthesis and used as template for qPCR using specific primer sets(42; 46). cDNA synthesis was performed on 5μg RNA per sample using Superscript first-strand cDNA kit (Invitrogen) and qPCR was performed by employing SYBR green supermix (Biorad). For bacterial DNA levels, total DNA was prepared from the fecal pellets(47; 48). qPCR assay was carried out using universal (total bacterial)- and BF- specific 16S rRNA gene targeted probe sets and SYBR green Supermix as described before(42).

### 16S rRNA Gene Sequencing and bacterial community profiling

Total DNA was extracted from the fecal pellets for bacterial community profiling as detailed in our previous reports(20; 43; 46-49). DNA in the samples was amplified by PCR using 16S rRNA gene-V3-V4 region targeted amplicon primer sets, and the sequencing was performed using Illumina MiSeq platform with the help of NC State University Genomic Sciences Laboratory (Raleigh, NC, USA) or Molecular Research LP (Shallowater, TX). The sequencing reads were fed into the Metagenomics application of BaseSpace (Illumina) for performing taxonomic classification of 16S rRNA gene amplicon reads using an Illumina-curated version of the GreenGenes taxonomic database, which provided raw classification output at multiple taxonomic levels. The sequences were also fed into QIIME open reference operational taxonomic units (OTU) picking pipeline (50) using the QIIME pre-processing application of BaseSpace. The OTUs were compiled into different taxonomical levels based upon the percentage identity to GreenGenes reference sequences (i.e. >97% identity), and the percentage values of sequences within each sample that map to respective taxonomical levels were calculated. The OTUs were also normalized and used for metagenomes prediction of the Kyoto Encyclopedia of Genes and Genomes (KEGG) orthologs by employing PICRUSt as described before(51–54). The predictions were summarized to multiple levels and functional categories as well as microbial communities at different taxonomical levels were compared among different groups. MicrobiomeAnalyst(55) and the Statistical Analysis of Metagenomic Profile Package (STAMP)(52) applications were employed for statistical analysis and visualization of data.

### Bulk RNA-seq analysis of gene expression

Bulk RNA-seq assay was done with the help of Novogene Inc using polyA mRNA isolated from the total RNA preparations sent to this service provider (detailed in supplemental method). Raw sequencing count data were then normalized and subjected to visualization and statistical analysis by employing web-based integrated Differential Expression & Pathway analysis (iDEP) 2.01 platform (https://bioinformatics.sdstate.edu/idep/(56)) as described in supplemental method.

### Adoptive cell transfer

In vitro β-cell antigen peptide cocktail stimulated or unstimulated pancreatic LN cells from 12- week-old mice were transferred into 8-week-old NOD-*Scid* mice (i.v.; 1 x 10^6^ cells/mouse) and tested for blood glucose levels every week as described above to determine the diabetogenic properties of T cells.

### Staining of intestinal tissue sections

Distal colon and ileum tissues were collected with luminal content, fixed overnight in formalin, 5μm sections were cut and used for immunofluorescence and histochemical staining. Tissue sections were stained using mouse Muc2 specific antibody (Novus Biologicals) followed by Alexaflor-488 conjugated secondary antibody. Tissue sections were also stained using Alcian blue/nuclear fast red, Periodic acid-Schiff’s reagent/Hematoxylin, and Alcian blue/Periodic acid-Schiff’s reagent/Hemotoxylin staining to detect mucin. All fluorescent and bright-field images were acquired using Evos 5000 microscope (Invitrogen).

### Statistical analysis

Mean, SD, and statistical significance (*p-value*) were calculated using GraphPad Prism, Microsoft Excel, and/or online statistical applications. Wilcoxon signed-rank test was employed, unless specified, for values from *in vitro* and *ex vivo* assays. Specific methods employed upon statistical analysis and visualization of RNAseq and 16s rRNA gene amplicon sequencing data are mentioned in figure legends. Log-rank analysis was performed to compare T1D incidence (hyperglycemia onset) of the test group with that of respective control group. Fisher’s exact test was used for comparing the total number of severely infiltrated islets (grades ≥3) relative to the total number of islets with low or no infiltration (grades ≤2) in test vs. control groups. A *p* value of ≤0.05 was considered statistically significant.

## Results

### Gut colonization of NOD mice by BF causes accelerated disease progression/hyperglycemia onset

Hyperglycemia is detected in female NOD mice housed in our facility as early as 12 weeks of age and 80-100% of the mice turn overtly hyperglycemic before the age of 30 weeks(20; 42; 43). Our previous report, which employed adult NOD mice, showed that systemic administration of small amount of HK BF accelerated the disease onset. Here, we used 15 to 16 -day-old NOD mice to determine the impact of oral administration of live BF for 3 consecutive days on eventual autoimmune progression and T1D incidence. The qPCR assay showed that live BF recipient mice had detectable levels of BF-specific 16S rRNA gene in the fecal samples collected 30 days post-colonization **(Fig. 1A)** suggesting successful gut colonization by this organism. As shown in **Fig. 1B**, gut colonization of NOD mice at juvenile age by BF caused a significant acceleration of hyperglycemia onset and overall higher T1D incidence during the 30-week monitoring period. To determine the impact of juvenile gut colonization by BF on islet autoimmunity, one set of mice was euthanized at 10 weeks of age, and the pancreatic tissues were examined for insulitis severity. Early onset and higher incidence of the disease in BF-colonized mice were found to correlate with significantly higher number of islets with more severe insulitis (grades ≥3 insulitis) **(Fig. 1C)**. Spleen cells from mice that received BF showed higher frequencies of IFNγ^+^ and IL6^+^ and lower frequencies of IL10^+^ CD4 cells **(Fig. 1D)**. Further, PnLN cells from BF- colonized mice, compared to control mice, not only produced higher amounts of pro-inflammatory cytokines IFNγ and IL6 upon ex vivo exposure to β-cell antigen peptides **(Fig. 1E)**, but also caused, upon adoptive transfer, relatively faster onset of hyperglycemia in NOD-*scid* mice **(Fig. 1F)**. Of note, depletion of gut microbes using broad spectrum antibiotics resulted in delayed onset of hyperglycemia in both control and BF-colonized mice **(Fig. 1G)**. Overall, these observations suggest that gut colonization by the human gut microbe BF at juvenile age causes a systemic pro-inflammatory response and promotes T1D disease progression in genetically susceptible mouse model.

**FIGURE 1:**
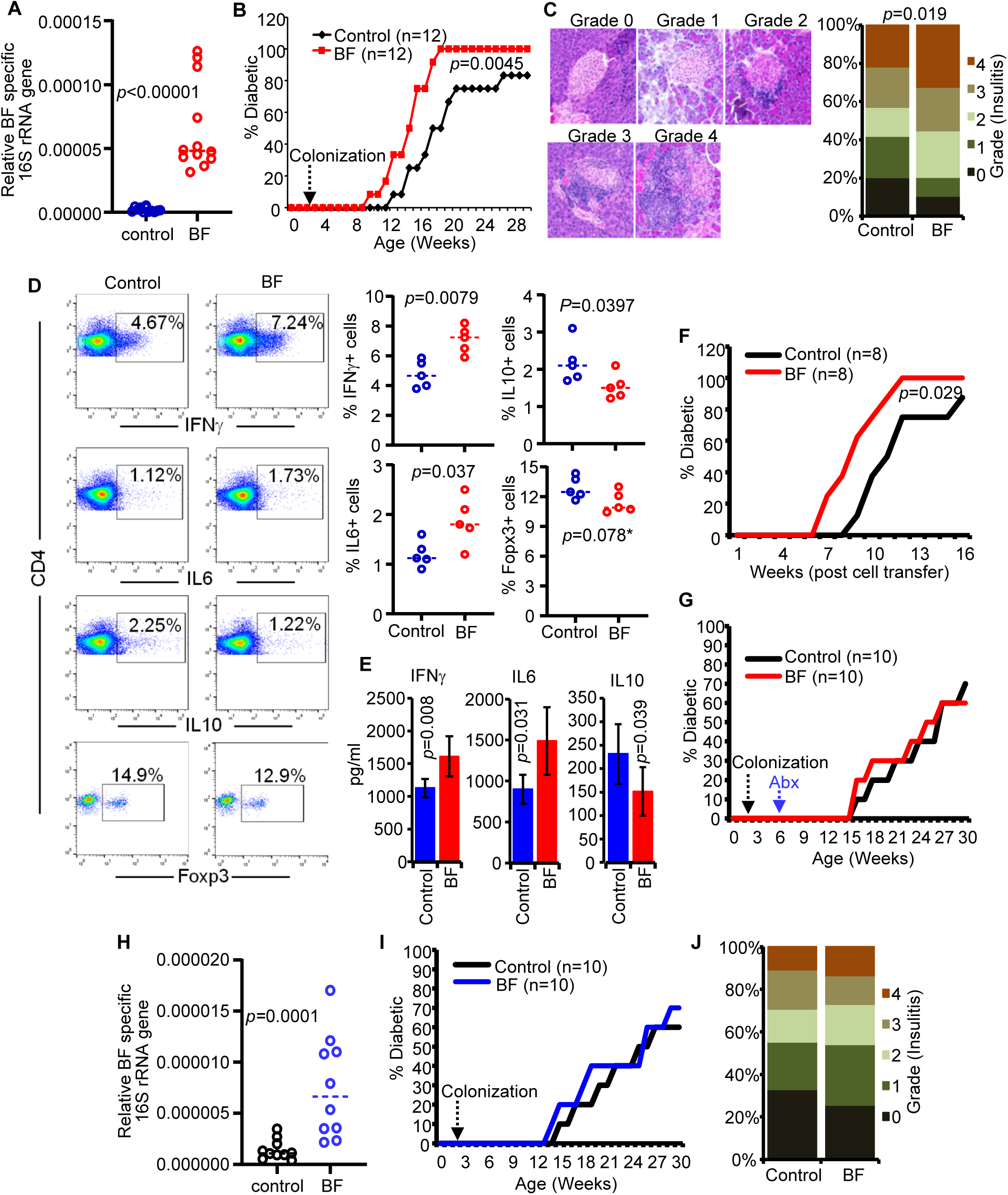
Accelerated disease progression and hyperglycemia onset, proinflammatory immune features in BF-colonized NOD mice. Juvenile (15-16-day-old) female NOD mice of conventional facility were given live BF (approx.:1x10^6^ CFU/mouse/day) or control broth for three consecutive days by oral gavage. **A)** DNA preparations of fecal samples collected after 30 days were tested for BF-specific 16S rRNA gene levels, relative to total bacterial 16S rRNA gene levels, by qPCR. **B)** Control and BF-colonized mice were monitored for hyperglycemia by weekly testing of blood glucose levels. Mice with glucose levels of >250mg/dl, for two consecutive weeks, were considered diabetic. Percentage of mice that developed diabetes at weeks post-treatment initiation are shown. **C)** Cohorts of mice (n=5/group) were euthanized at 10 weeks of age, pancreatic tissue sections were subjected to H&E staining and examined for insulitis. Examples of islets with different insulitis grades (left panel) and the percentage islets with different insulitis grades (right panel) are shown. **D)** Spleen cells were subjected to intracellular/intranuclear staining for indicated factors and analyzed by FACS. Representative FACS graphs (left) and percentage of CD4+ cells that are positive for specific markers (right) are shown. **E)** PnLN cells were cultured for 24h in the presence of β-cell antigen peptide cocktail and supernatants were tested for the indicated cytokine levels by multiplex assay. **F)** Peptide activated PnLN cells from 24h cultures were injected (i.v.) into NOD-*Scid* mice (Approx: 1x10^6^ cells/mouse) and monitored for hyperglycemia. **G)** At 6 weeks of age, control and BF-colonized mice were given drinking water containing broad spectrum antibiotic cocktail for 15days and monitored for hyperglycemia. Juvenile NOD-TLR2^-/-^ mice were given live BF as described above. **H)** Fecal samples were tested for BF-specific 16S rRNA gene levels as described for panel A. **I)** Cohorts of mice were monitored for hyperglycemia as described for panel B. **J)** Pancreatic tissues from cohorts of mice (n=5) were examined for insulitis grades as described for panel C. Statistical significance by Mann-Whitney test for panels A, D, E and H; log-rank test for panels B and F; and Chi-square test for panel C. *not significant.

Our previous report(42), using TLR2 deficient NOD mice, showed that HK BF induced acceleration of T1D progression is TLR2 dependent. Therefore, we examined if juvenile age BF colonization induced disease acceleration effects on T1D in NOD mice are TLR2 dependent. Juvenile (15-16 day-old) female NOD-TLR2-KO mice were given live BF and examined for colonization efficiency **(Fig. 1H)**. Cohorts of BF-colonized and control NOD-TLR2-KO mice were monitored for hyperglycemia onset for 30 weeks or euthanized at 10 weeks of age and the pancreatic tissues were examined for insulitis. As observed in **Fig. 1I-1J**, unlike WT NOD mice (Fig. 1B), colonization by BF did not produce significant modulatory effects on disease progression and insulitis in TLR2-KO mice compared to control mice. These results confirm our previous report(42) on a role for BF-TLR2 interaction in BF induced systemic pro-inflammatory response and T1D promoting effects in NOD mice.

### Gut microbiota composition is significantly different in BF-colonized and control NOD mice

Since depletion of gut microbiota eliminated BF-colonization-associated acceleration of T1D disease progression, we compared the microbial community profiles of BF-colonized and control NOD mice. Fecal samples collected 4 weeks post-BF-colonization were subjected to 16s rRNA gene amplicon sequencing and analyzed for overall diversity and composition of microbiota. Principal component/β-diversity analysis of the sequence data showed significantly distant clustering of samples from control and BF-colonized mice **(Fig. 2A)** suggesting that the gut microbial community structure is influenced by BF. Importantly, BF-colonized mice showed relatively higher microbial diversity compared to their control counterparts as indicated by differences in the α-diversity measures/Chao1 index **(Fig. 2B)**. Sequence data were also analyzed to determine the abundance of microbial communities at phyla and genus levels. Among major phyla, the abundance of Bacteroidetes members was significantly higher and Firmicutes was lower in BF-colonized mice compared to controls **(Fig. 2C)**. At the genus level, lower abundance of *Lactobacillus* and higher abundance of *Bacteroides* were observed in BF- colonized mice compared to control counterparts **(Fig. 2D)**. These results suggest that BF colonization early in life could, in addition to TLR2-dependent effect, alter the gut microbiota composition and impact disease progression in T1D.

**Figure 2:**
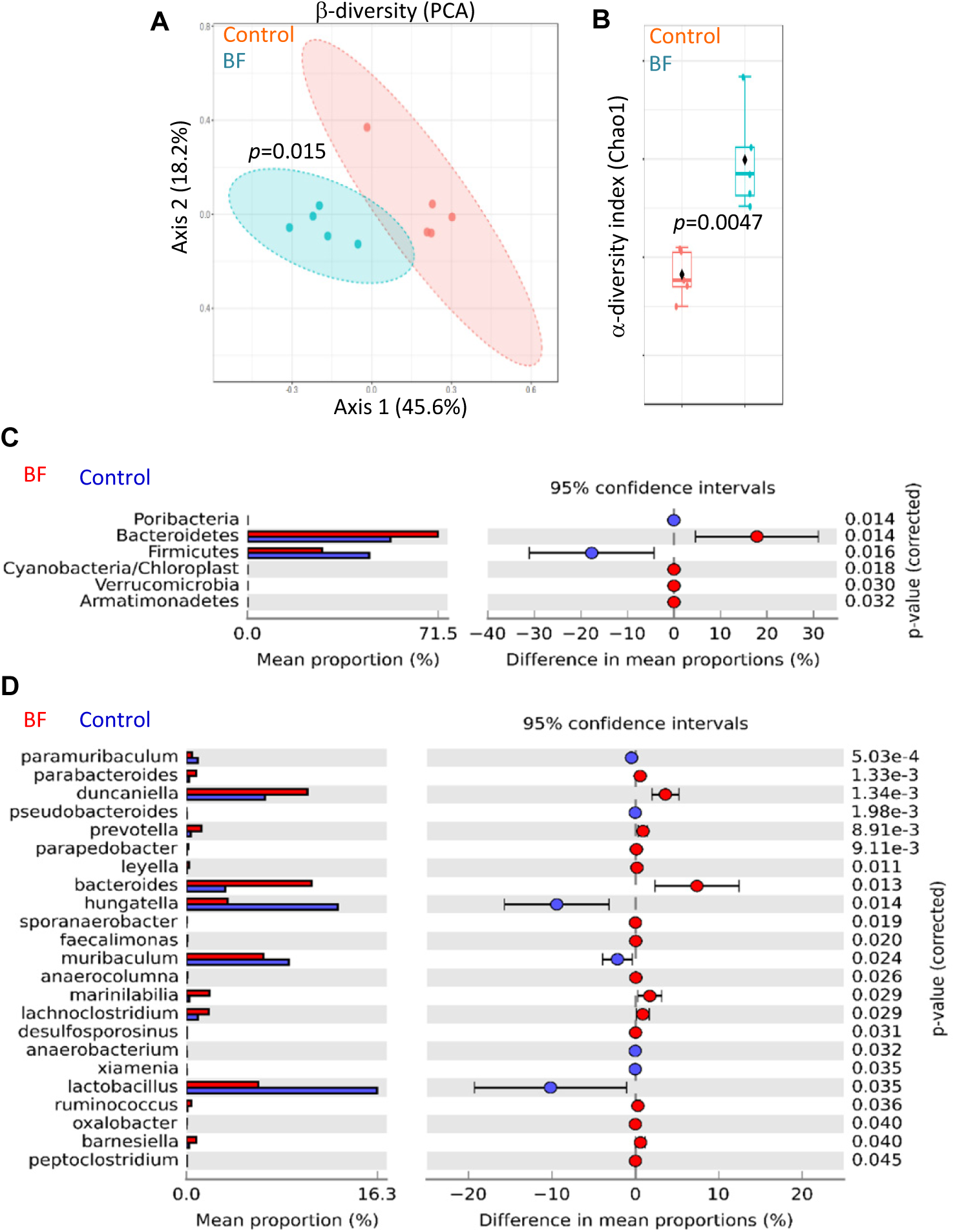
Differences in the fecal microbiota composition in BF-colonized and control NOD mice. Fresh fecal pellets were collected from control and BF-colonized mice at 6 weeks of age, DNA preparations were subjected to 16S rRNA gene (V3/V4 region) -targeted sequencing using the Illumina MiSeq platform. The OTUs that were compiled to different taxonomical level based upon the percentage identity to reference sequences (i.e. >97% identity) and the percentage values of sequences within each sample that map to specific phylum and genus were calculated by employing 16S Metagenomics application of Illumina BaseSpace hub. These sequencing data were subjected to visualization and statistical analyses employing Microbiomeanalyst or STAMP applications. **A)** Principal component analysis (PCA) plots of samples representing β-diversity (Bray Curtis distance). **B)** α-diversity (Chao1) comparison of control and BF groups. **C)** Mean relative abundances of microbial communities at phyla level. **D)** Mean relative abundances of sequences representing microbial communities at genus level. Statistical analyses were done employing two-sided Welch’s t-test and the *p*-values were FDR corrected using Benjamini and Hochberg approach for panel D.

### Monocolonization of GF NOD mice by BF results in accelerated disease progression

Since the gut microbiota composition is different in control and BF-colonized mice, the impact of gut colonization of NOD mice by BF, in the absence of other microbes, on disease progression was examined. GF NOD mouse breeders in gnotobiotic isolators were monocolonized with BF and the female litters were tested for BF specific DNA levels in their fecal samples **(Fig. 3A)**. These mice, along with age matched GF NOD mice were monitored for hyperglycemia. As observed in **Fig. 3B**, BF-monocolonized mice showed early onset and rapid progression to hyperglycemia compared to GF NOD mice. Examination of pancreatic tissues from a cohort of mice showed significantly higher frequencies of islets with more severe immune cell infiltration in BF-monocolonized mice compared to GF mice. Spleen cells from BF-monocolonized mice produced profoundly higher amounts of pro-inflammatory cytokines including IL6, IL12, TNFα, IFNγ, IL17, IL18, IL13, GM-CSF and IL1β compared to their GF counterparts **(Fig. 3C)**. qPCR assay showed that distal colon tissues from BF-monocolonized mice expressed higher levels of mRNA for pro-inflammatory cytokines such as TNFα, IFNγ, IL6, IL12, and IL1β and lower levels of IL10 and Ido1 **(Fig. 3D)**. Importantly, PnLN cells from BF-monocolonized mice showed higher frequencies of IFNγ^+^ and lower frequencies of Foxp3+ CD4 cells compared to GF control mice **(Fig. 3F)**. Further, adoptive transferred PnLN cells from BF-monocolonized NOD mice, but not the cells of GF NOD mice, to NOD-*Scid* mice caused hyperglycemia **(Fig. 3G)**. Overall, these results show that gut colonization by BF early on in life can cause a pro-inflammatory immune response and accelerate autoimmune progression in T1D-prone/genetically susceptible mice.

**Figure 3:**
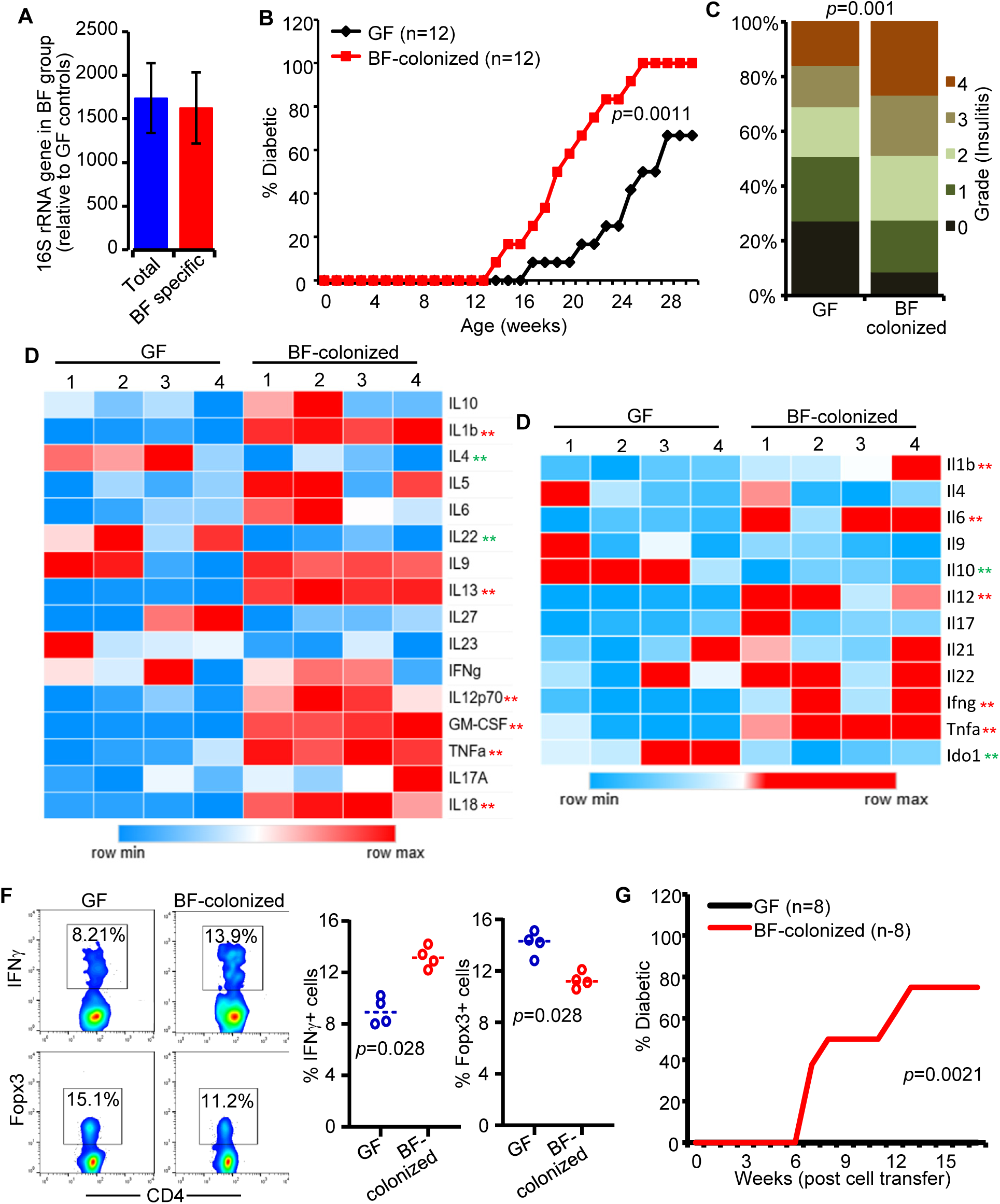
Accelerated T1D disease progression and pro-inflammatory immune response in BF-monocolonized NOD mice. A) Fecal samples from 5 randomly selected female litters of BF-monocolonized NOD mouse breeders in the gnotobiotic facility were tested for total bacterial and BF-specific 16S rRNA gene levels relative to the background values of fecal samples from their GF counterparts by qPCR. **B)** Cohorts of GF control and BF-monocolonized mice were monitored for hyperglycemia for up to 30 weeks of age as described under Materials and methods. **C)** Pancreatic tissue sections of cohorts of 12-week-old mice (n=4/group) were subjected to H&E staining and examined for insulitis grades. **D)** Spleen cells from 12-week-old mice were cultured in the presence of anti-CD3 antibody for 24h and supernatants were tested for cytokine levels by Luminex multiplex assay. **E)** RNA prepared from the distal colon tissues was subjected to qPCR to determine the expression levels of indicated factors. **F)** PnLN cells from a cohort of 12-week-old mice were stained for intracellular/intranuclear markers and analyzed by FACS. Representative FACS graphs (left) and percentage of IFNγ+ and Foxp3+ cells among CD4+ cells (right) are shown. **G)** Pooled, unstimulated, PnLN cells from a similar cohort of mice were injected in to 8–10-week-old NOD-*Scid* mice (1x10^6^ cells/mouse) and monitored for diabetes as described above. Statistical significance by log-rank test for panels B and G; Chi-square test for panel C; Mann-Whitney test for panels D, E and F. Statistically significant **upregulation or **down regulation in BF-monocolonized mice compared to GF mice.

### BF-colonization alters gene expression profiles in the distal gut of mice under gnotobiotic condition

Although qPCR analysis of BF-monocolonized mice showed differences in the immune characteristics of distal colon compared to GF controls (Fig. 3), the impact of colonization by BF, in the absence of other microbes, on distal gut gene expression profile was assessed further by employing bulk-RNAseq. Differential gene expression (DEG) profiles of BF-monocolonized vs GF control NOD mice were examined. BF-monocolonization alone caused major differences in the overall gene expression profile **(Fig. 4A)** and changes in the expression levels of a large number of genes, 1149 upregulated and 437 downregulated **(Fig. 4B-C)**, in the distal colon. The prominent genes upregulated upon BF monocolonization include *Acta2*, *Tpm1*, *Tpm2*, *Cald1*, *Mylk*, *Des*, *Myl9*, and *Tns1*, which are involved in smooth muscle and fibroblast functions. On the other hand, major genes that are downregulated include *Slfn4*, which is involved in myelopoiesis and macrophage function, and *Gasdmc2* and *Gasdmc3* that contribute to pyroptosis, epithelial cell function, and host defense to enteric pathogens **(Fig. 4D)**. Pathway enrichment analysis using gene lists of DEG by KEGG approach confirms modulation of these cellular functions in the distal gut of BF-monocolonized mice **(Fig. 4E**). On the other hand, pathway analysis done using fold change values of all genes by GO approach showed down regulation of IL10 production pathway including lower *Cd274* (PD-L1), *Pdcd1lg2* (PD-L2) and *Ido1* expression **(Fig. 4F)**. In fact, this is consistent with our qPCR results showing diminished expression of *Il10* and *Ido1* in the distal gut of BF-monocolonized mice **(Fig. 3)**. Although the sequence counts of many of the immunological factors were extremely low, low-count sequence data revealed that, consistent with qPCR data **(Fig. 3)**, the expression levels of many of the pro-inflammatory cytokines were upregulated in BF-monocolonized mice compared to that of their GF counterparts (not shown). Overall, these observations in association with the systemic immune response of BF-monocolonized mice **(Fig. 3)** explain the rapid T1D disease progression and hyperglycemia onset in BF-colonized mice.

**Figure 4:**
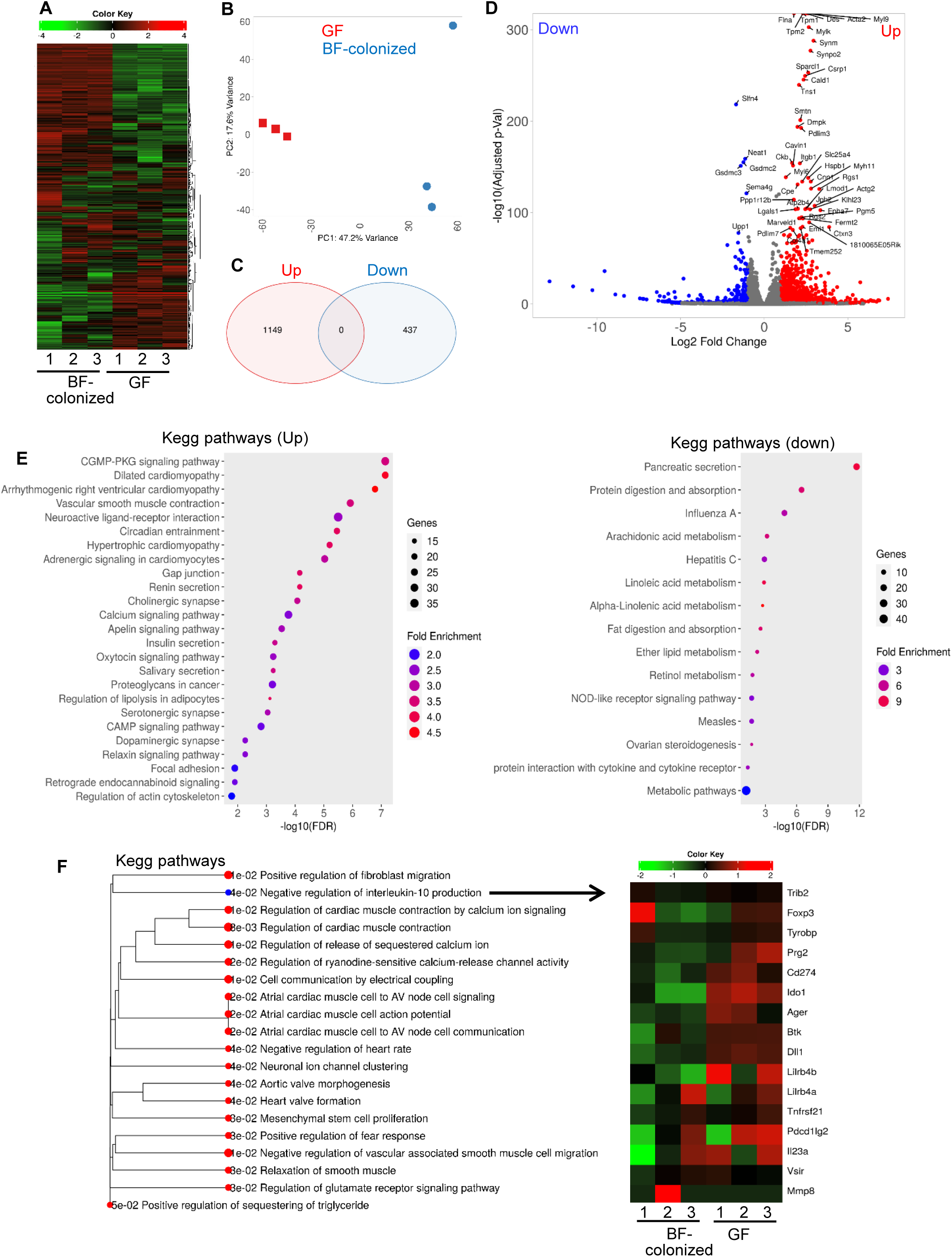
Differences in gene expression profiles of the distal gut from GF and BF- monocolonized NOD mice. RNA prepared from the distal colon of 12-week-old GF control mice and BF-monocolonized mice were subjected to bulk-RNAseq and the gene counts of data were then normalized and subjected to visualization and statistical analysis using iDEP2.01 platform. **A)** Heatmap showing hierarchical clustering, employing Pearson’s correlation coefficient, of top 2000 gens showing holistic view of differences in the sequencing data of individual samples. **B)** PCA graph showing overall differences in the gene expression profiles among samples and groups. **C)** Venn diagrams showing number of significantly up- and down-regulated genes among DEGs (identified employing DESeq2 with a minimum fold change of 2 and FDR cutoff as 0.05) in BF-colonized mice compared to GF control mice. **D)** Volcano plot showing significantly up- and down-regulated genes among DEGs in BF-colonized mice compared to GF mice with top 50 genes labeled. **E)** KEGG Pathway enrichment analysis of significantly up- and down-regulated metabolic pathways or signal transduction pathways associated with DEG. **F)** KEGG pathway enrichment analysis also used for fold-change values of all genes, independent of the selected DEG, to identify up- and down-regulated pathway, and the differences in expression profiles of genes associated with negative regulation of IL10 production are shown.

### Acceleration of T1D disease process persisted in BF-monocolonized mice even after conventionalization

To examine whether introduction of conventional facility microbiota has an impact on BF monocolonization-associated acceleration of disease progression, GF and BF-monocolonized NOD mice were pulled from the gnotobiotic isolators immediately after weaning (about 4 weeks of age) and housed (conventionalized) on the pooled and evenly distributed dirty bedding from NOD mouse breeding cages of conventional facility. Fecal microbial levels were assessed using samples collected after 1 month of conventionalization by anaerobic and aerobic BHI agar plate cultures (not shown). BF specific 16s rRNA gene levels in the fecal samples were also determined at different time-points by qPCR assay to assess continued presence of BF **(Fig. 5A)**. Cohorts of conventionalized mice were monitored for blood glucose levels to determine T1D incidence. As observed in **Fig. 5B**, although conventionalization alone caused a delay in hyperglycemia onset compared to mice of conventional and gnotobiotic facilities (Figs.1&3), conventionalized BF-monocolonized mice (Ex-BF / BF-conv mice) showed higher incidence of diabetes during the monitoring period of up to 30 weeks of age compared to conventionalized GF mice (Ex-GF / GF-conv mice). Relatively higher insulitis severity was observed in pancreatic tissues of Ex-BF mice, collected from a cohort of mice at 8 weeks post-conventionalization compared to Ex-GF mice **(Fig. 5C)**. Cytokine secretion profiles of spleen cells showed considerable differences, with the Ex-BF mice presenting overall higher proinflammatory immune response compared to the Ex-GF mice **(Fig. 5D)**. Similar pro-inflammatory features were observed with the distal colon tissues from Ex-BF (not shown). Overall, these results demonstrate that BF colonization early in life can produce lasting impact on pathogenesis in T1D even in the presence of diverse gut microbial communities.

**Figure 5:**
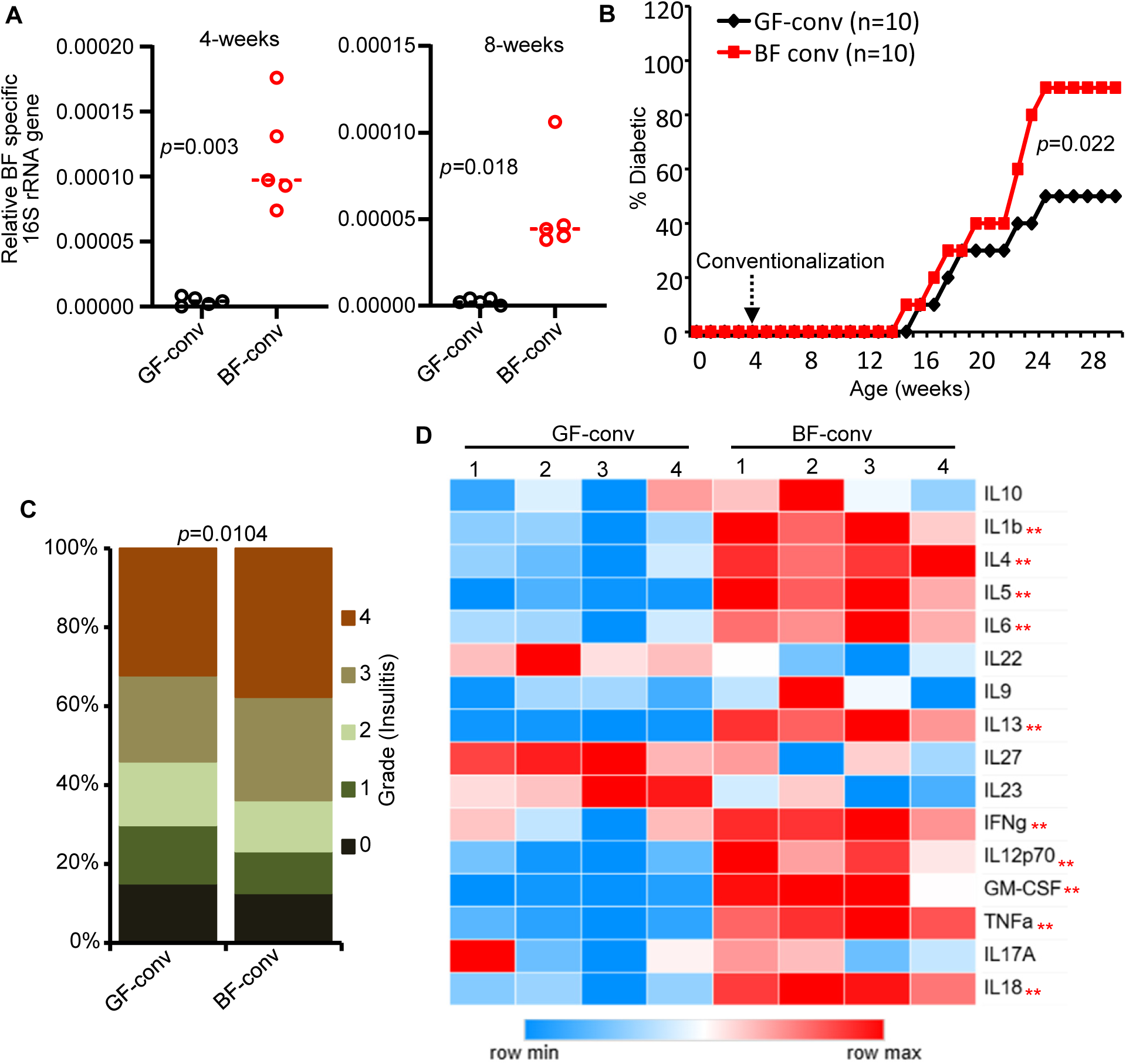
Accelerated T1D disease process in conventionalized BF-monocolonized mice. Four-week-old GF mice and BF-monocolonized -NOD mice were housed on pooled and equally distributed dirty bedding of conventional NOD mice. Fecal samples from randomly selected conventionalized GF (GF-conv / Ex-GF) and conventionalized BF-monocolonized (BF-conv / Ex-BF) mice (n=5) were examined for BF-specific 16S rDNA gene levels relative to total 16S rDNA gene levels by qPCR assay. **B)** Cohorts of conventionalized mice were monitored for hyperglycemia for up to 30 weeks of age. **C)** Cohorts of euglycemic mice (n=5/group) were euthanized at 12 weeks of age, pancreatic tissue sections were subjected to H&E staining and examined for insulitis grades. **D)** Spleen cells were cultured in the presence of anti-CD3 antibody for 24h and supernatants were tested for cytokine levels by Luminex multiplex assay. Statistical significance by Mann-Whitney Chi-square test for panels A and D, log-rank test for panels B and Chi-square test for panel C. Statistically significant **upregulation or **down regulation in Ex-BF mice compared to Ex-GF mice.

### Presence of BF impacts acquisition and overall structure and function of gut microbiota

To determine if presence of BF has an impact on the acquisition and persistence of gut microbial communities, fecal samples collected from a cohort of Ex-GF control and Ex-BF groups of mice at 4- and 8-weeks post-conventionalization were subjected to 16s rRNA gene amplicon sequencing and analyzed to determine the microbial community profiles. Significant differences in the α- and β-diversity measures were observed between Ex-GF and Ex-BF mice with higher microbial diversity in Ex-BF mice at both 4- and 8-week time-points **(Fig. 6A)**. Among major phyla, the overall abundances of Bacteroidetes members are significantly higher and Verucomicrobia and Proteobacteria members are profoundly lower in EX-BF mice at both 4- and 8-week timepoints after conventionalization **(Fig. S1)**. Since both groups of mice were conventionalized using the dirty bedding from a single source, we compared the microbial community profiles down to genus and species level. As shown in **Fig. 6Band S2**, while *Akkermansia muciniphila* and *Parabacteroides distasonis* were less abundant in Ex-BF mice, *Muribaculum intestanale* and *Barnesiella intestinihominis of Barnesiella genus* and Bacteroides genus members showed higher abundance in these mice compared to Ex-GF controls. When compared to Ex-GF mice, decrease in *A. muciniphila* in Ex-BF mice was more pronounced than any other microbial species at both time-points, with near complete absence of this species in Ex-BF mice at the 8-week timepoint. We then employed PICRUSt application for in silico prediction of metabolic functions that may be associated with differences in gut physiology in the presence and absence of BF. **Fig. S3** shows that a variety of microbial metabolic pathways including glycan biosynthesis and metabolism and energy metabolism pathways are overrepresented and lipid, xenobiotic and carbohydrate metabolism pathways are underrepresented in Ex-BF mice compared to Ex-GF control mice. Although further analysis and additional studies are needed to understand if such metabolic pathways of gut microbiota contribute to modulation of intestinal and systemic immune responses and T1D disease progression, these observations suggest that microbial community structure and function are different in the presence and absence of BF. Furthermore, this data suggests that presence of BF in the gut alone can have a profound influence on acquisition and maintenance of specific microbial communities.

**Figure 6:**
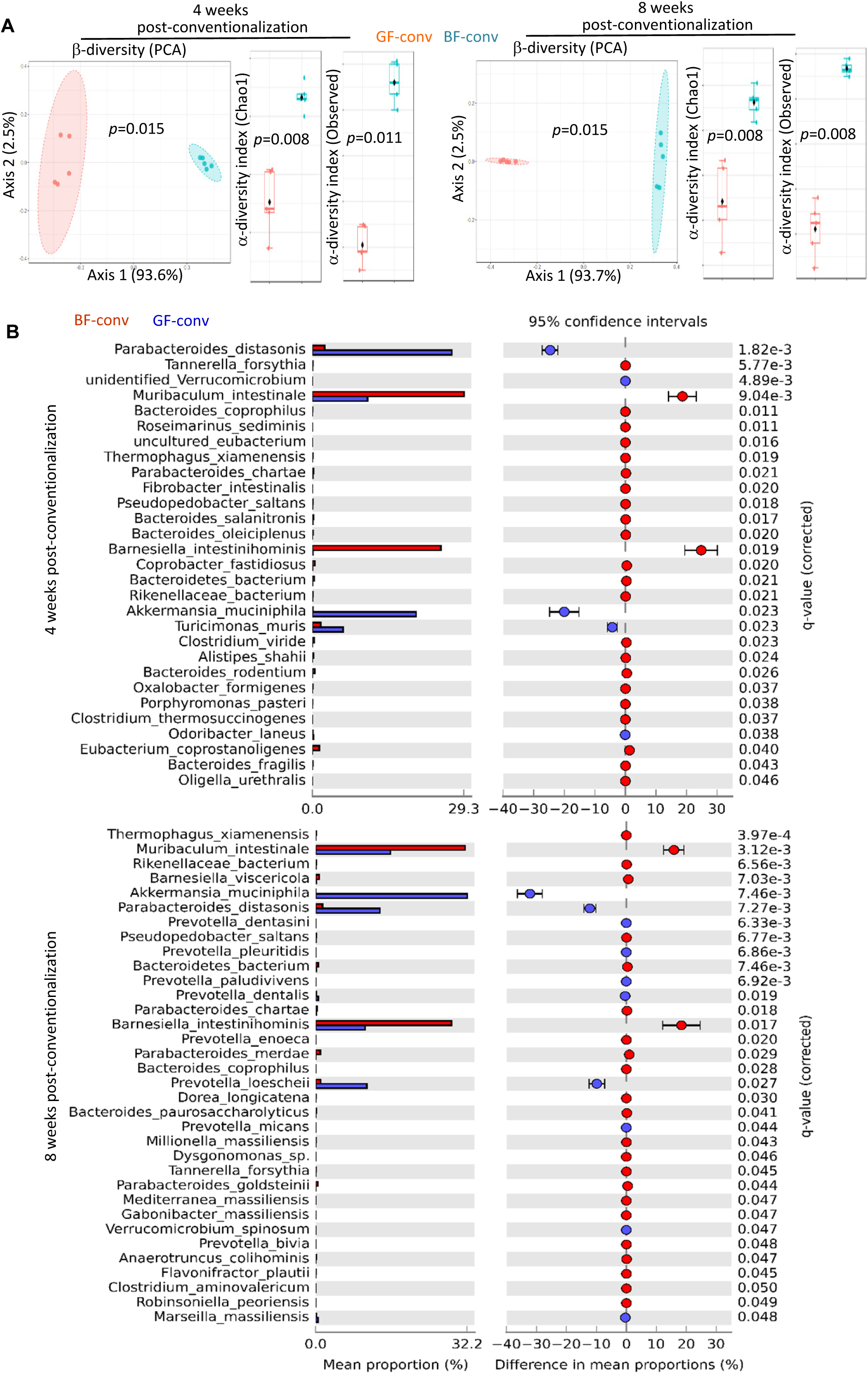
**Impact of BF on the acquisition and overall community profiles of gut microbiota**. Fecal pellets were collected from conventionalized GF (GF-conv / Ex-GF) and conventionalized BF-monocolonized (BF-conv / Ex-BF) mice at 4- and 8-weeks post-conventionalization, and the DNA preparations were subjected to 16S rRNA gene (V3/V4 region) -targeted sequencing using the Illumina MiSeq platform and the sequence data was analyzed as described for Fig. 2A) PCA plots representing β-diversity (Bray Curtis distance) and α-diversity (Chao1 or observed species) comparison. **B)** Mean relative abundances of sequences representing microbial communities at species level. Statistical analysis: two-sided Welch’s t-test and the *p*-values were FDR corrected using Benjamini and Hochberg approach.

### Presence of BF impacts gene expression profiles of distal gut in conventionalized NOD mice

Since Ex-GF and Ex-BF mice showed significant differences in the gut microbiota, disease incidence rate, and insulitis grades, gene expression profiles of distal colon tissues from these mice were examined by bulk RNAseq. Heatmap and PCA plots of RNAseq data showed differences in the expression profiles and distant clustering of Ex-GF group samples from Ex-BF group samples **(Fig. 7A and 7B)** suggesting major differences in the gene expression profiles of these groups. DEG analysis showed that presence of BF in conventionalized NOD mice resulted in the upregulation of 523 genes and downregulation of 250 genes **(Fig. 7C and 7D)**. DEG analysis also showed that some of the major upregulated genes included *SLC37A2* and *Atp12a* with membrane channel functions and ATP12A with H+/K+-ATPase function, mucin gene MUC2 and mucin synthesis associated gene *Gcnt3* and tight junction protein gene *Cldn15* expressions were down regulated in the distal colon of Ex-BF mice **(Fig. 7E)**. KEGG pathway enrichment analysis employing DEG profiles revealed upregulation of many immunological pathways, including those related to Th1, Th2 and Th17 differentiation, B cell receptor signaling and IgA production, and autoimmune and inflammatory bowel diseases **(Fig. 7F)**. On the other hand, presence of BF in the conventionalized mice caused downregulation of mucin biosynthesis pathway along with profoundly diminished expression of *Muc2* and other mucin genes. GO enrichment analysis for biological processes employing DEG list also showed upregulation of several immune function associated processes and down regulation of glycoprotein biosynthesis, glycosylation, and glycoprotein metabolic processes **(Fig. 7G)**. These results show that presence of BF results in suppression of mucin production associated genes and upregulation of proinflammatory immune pathway genes.

**Figure 7:**
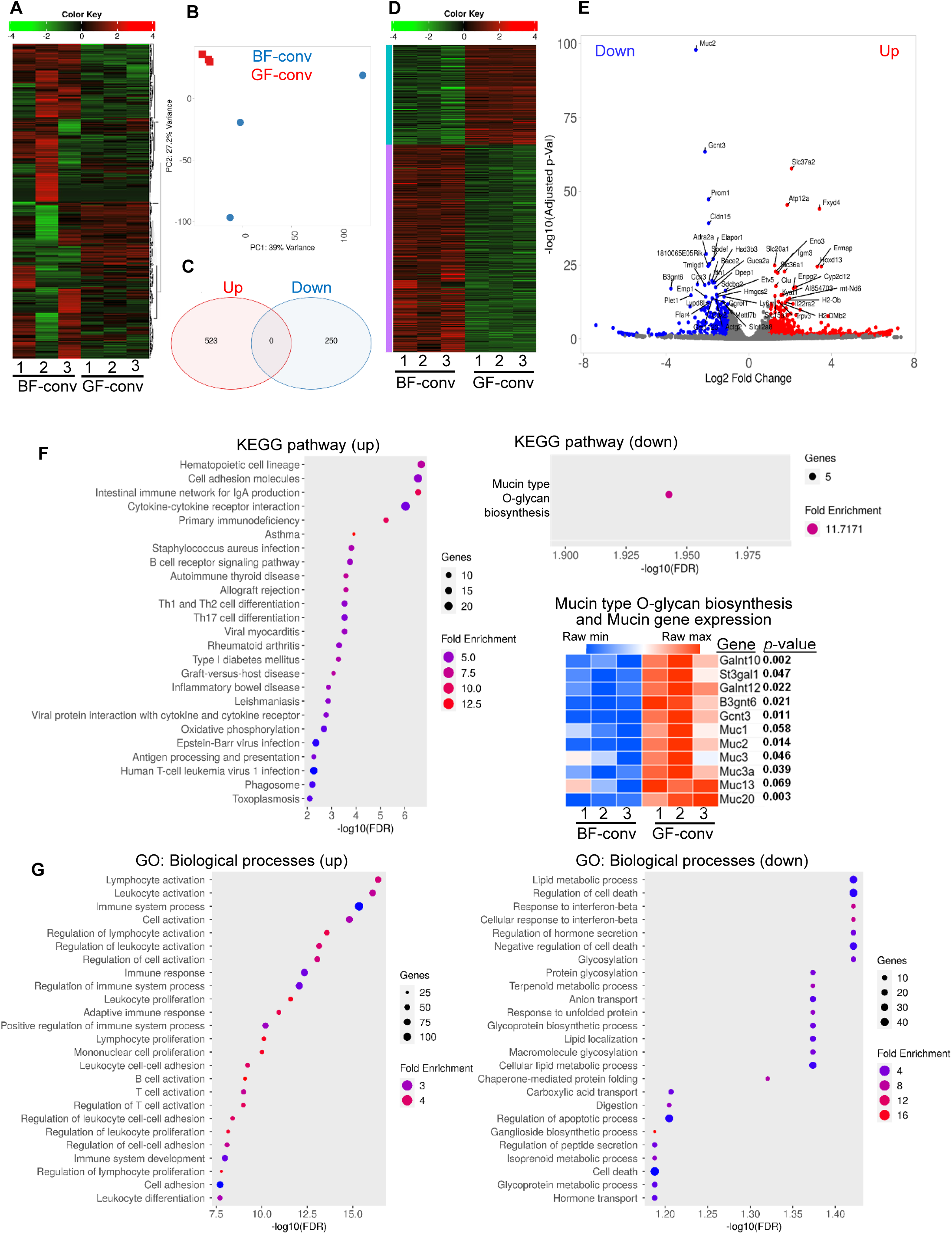
**Differences in the gene expression profiles of the distal gut from conventionalized GF and BF-monocolonized NOD mice**. RNA prepared from the distal colon of 12-week-old conventionalized GF (GF-conv / Ex-GF) and conventionalized BF-monocolonized (BF-conv/ Ex-BF) mice were subjected to bulk-RNAseq and analyzed as described for Fig. 4. **A)** Heatmap showing hierarchical clustering, employing Pearson’s correlation coefficient, of top 2000 genes showing differences in the sequencing data of individual samples. **B)** PCA graph showing overall differences in the gene expression profiles among samples and groups. **C)** Venn diagrams showing significantly up- and down-regulated genes among DEGs (identified employing DESeq2 with a minimum fold change of 2 and FDR cutoff as 0.05) in Ex-BF mice compared to Ex-GF control mice. **D)** Heatmap representing all DEGs in individual samples. **E)** Volcano plot showing significantly up- and down-regulated genes among DEGs in Ex-BF mice compared to Ex-GF mice with top 50 genes labeled. **F)** KEGG Pathway enrichment analysis of significantly up- and down-regulated metabolic pathways or signal transduction pathways in Ex-BF mice using DEG list, and heatmap comparing sequence counts of mucin biosynthesis associated and mucin genes. **G)** GO enrichment analysis of significantly up- and down-regulated biological processes in Ex-BF mice using DEG list.

### Presence of BF results in suppressed mucin production in the distal gut epithelium

Since the expression levels of Muc2 and mucin synthesis associated genes were found to be down regulated in BF-colonized mice, distal colon tissues sections of a cohort of 12-week-old Ex-GF and Ex-BF NOD mice were examined for Muc2 protein expression by immunofluorescence assay and overall mucin levels by employing Alcian blue and Periodic Acid-Schiff’s (PAS) staining approaches. As observed in **Fig. 8A**, anti-Muc2 antibody staining revealed considerably weaker staining of Muc2+ cells in Ex-BF mice compared to Ex-GF NOD mice. Interestingly, staining intensity was profoundly weaker close to the crypt areas of distal gut epithelium. Of note, ileum sections from the same animals showed no considerable difference in the staining pattern for Muc2, suggesting that Muc2 expression is affected primarily in the BF colonization niche. Examination of mucin levels by Alcian blue and PAS staining also showed similarly lower mucin positivity in the distal colon and comparable staining intensities in the ileum of Ex-BF mice compared to Ex-GF mice **(Fig. 8B)**. Overall, these results in the context of Figs. 6 and 7 demonstrate that human gut commensal BF can cause remodeling of distal colon niche by inducing proinflammatory immune response and lowering mucin production, leading to suppression of colonization by specific mucin degrading microbial communities such as *Akkermansia muciniphila* and accelerated autoimmune progression in T1D.

**Figure 8:**
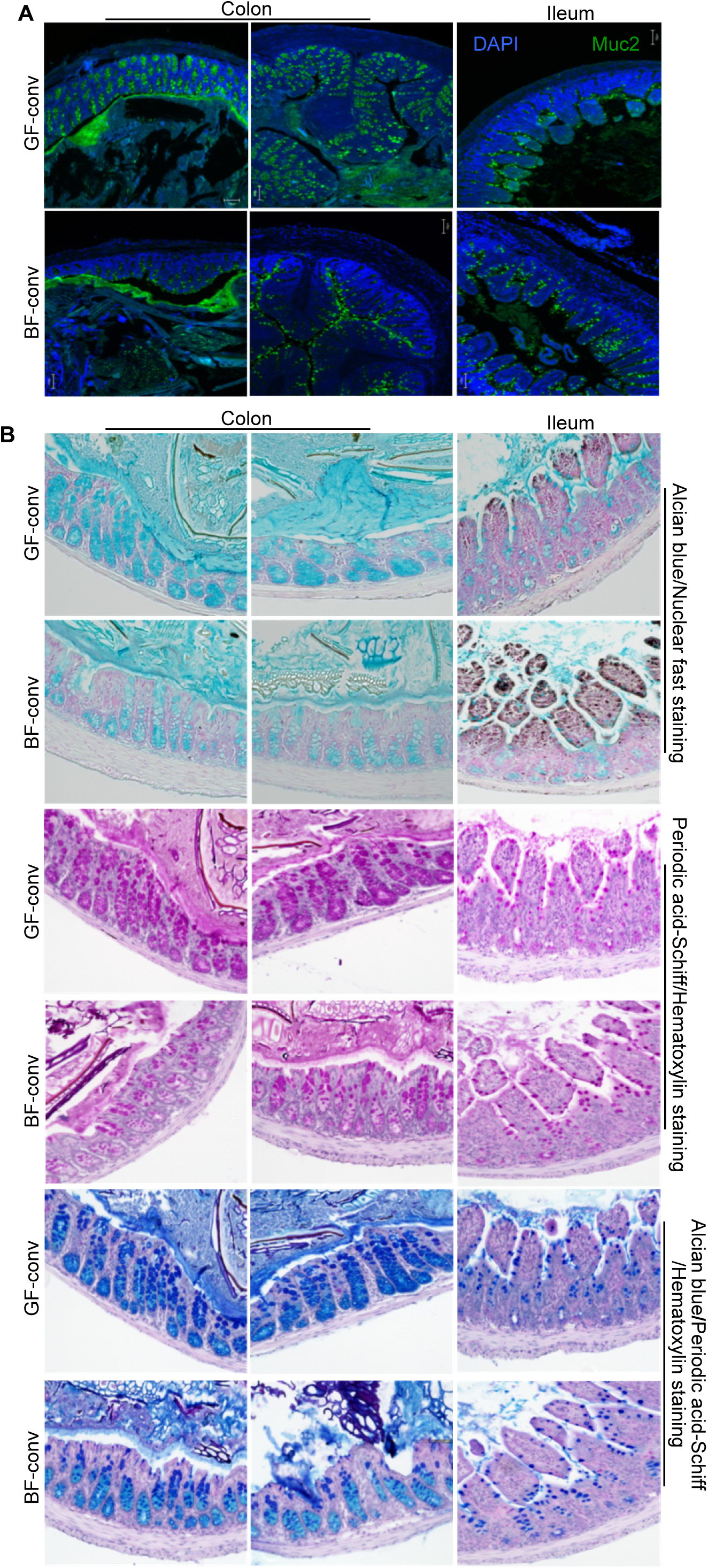
Suppressed mucin production in the distal gut of conventionalized BF-colonized mice. Sections of formalin fixed distal colon and ileum tissues (with luminal contents) of 12-week-old conventionalized GF (GF-conv / Ex-GF) and conventionalized BF-monocolonized (BF-conv / Ex-BF) mice were subjected to immunofluorescence staining using anti-Muc2 antibody **(A)**, or Alcian blue/nuclear fast, periodic acid-Schiff’s/hematoxylin, and Alcian blue periodic acid-Schiff’s/hematoxylin staining **(B)**. Note: Colon and ileum tissues from 3 mice/group were embedded in the same block, sectioned, and stained to minimize tissue-to-tissue variability in standing intensity and representative images are shown.

## Discussion

Gut commensal microbes are critical to human health and immune system maturation(1–4). However, reports also suggest that gut commensals can contribute to the pathogenesis of many autoimmune diseases including type 1 diabetes (T1D)(5–7). Studies have demonstrated the unique immune-regulatory properties and symbiotic nature of BF(27; 33). BF is also the primary microbe detected in most clinical intra-abdominal abscesses(25; 26; 57; 58), suggesting that it may promote immune regulation when restricted to the gut, but its escape to the systemic compartment could produce a pro-inflammatory response and promote autoimmune progression. Earlier, we tested this notion in a NOD mouse model of T1D, using HK BF and employing oral and i.v. administrations to mimic gut mucosa and systemic exposures to this organism(42). We showed that, in contrast to the exposure of gut mucosa to BF which produced protection from T1D, exposure of the systemic compartment to BF resulted in a pro-inflammatory response, rapid insulitis progression and early onset of hyperglycemia. Using a PSA deficient mutant of BF and NOD-TLR2 deficient mice, we also showed that these opposing effects of BF on T1D, upon oral and systemic administrations, are in part bacterial PSA and host TLR2 interaction dependent.

Higher abundance of Bacteroidetes phyla members and a reduction in Firmicutes are associated with T1D(16; 17; 20). However, protection from T1D has also been positively correlated with higher abundance of multiple species belonging to Bacteroidetes phylum, particularly *Prevotella* sp(17; 18). BF, which colonizes the colonic crypt, is known to promote gut immune regulation(27; 29). Our study employing HK BF showed that it can promote T1D pathogenesis, especially upon reaching the systemic compartment(42). While our previous study that used HK BF in NOD mice has tested this notion(42), whether gut colonization by live BF impacts T1D progression, especially at juvenile age, is unclear. Here, using conventional and GF NOD mice, we show that gut exposure of NOD mice to live BF at juvenile age causes a pro-inflammatory systemic immune response and accelerated T1D disease progression. Although adult age exposure of NOD mice in the conventional facility to BF produces modest protection from the disease (not shown), our overall observations suggest that juvenile age acquisition of BF-like organisms can produce adverse effects in T1D prone subjects. Interestingly, not only did the litters of BF-monocolonized NOD mice and BF-exposed mice of the conventional facility show accelerated disease outcomes, but conventionalization of BF-monocolonized mice did not fully reverse their disease progression trajectory, suggesting that early life (and potentially continued) exposure to BF-like microbes has a significant effect on the eventual disease outcomes.

Our previous report(42) and the observations of live BF in WT and TLR2 deficient mice showed that the juvenile-age colonization-induced effect is, in part, TLR2 dependent. However, microbial community profiling of BF-colonized mice from the conventional facility and the conventionalized BF-monocolonized mice suggests that BF colonization associated effects could also be influenced by continued presence and/or its impact on the overall structure and function of microbiota. Gut colonization by BF appears to have a profound impact on the acquisition of specific gut microbiota communities. The presence of BF appears to permit the acquisition of higher levels of microbial species such as *Muribaculum intestanale* and *Barnesiella intestinihominis* (both belongs to the order *Bacteroidales*) and restrict the colonization of other communities including *Akkermansia muciniphila, Parabacteroides distasonis and Prevotella* spp. Considering the fact that BF as well as *A. muciniphila, P. distasonis* and *Prevotella* spp. are mucin binding or utilizing bacteria(59–62), these observations are particularly of interest and suggest a potential competition among these microbes within the colonic niche, resulting in restriction and/or gradual elimination of other mucin degraders by BF. Previous reports showing the ability of BF and certain toxin-producing *Bacteroides* spp. to kill diverse *Bacteroides*, *Prevotella*, and *Parabacteroides* spp. (41; 63-65) also support the notion of their competitive interactions.

Our transcriptome data of distal colon tissues from Ex-BF and Ex-GF mice discovered more evidence of these potential competitive interactions. The observation that mucin-type O-glycan biosynthesis pathway and the expression of different mucin genes, including the primary mucin gene of gut mucosa *Muc2*, and the overall mucin levels are profoundly suppressed in the distal gut of BF-colonized mice, suggesting the possibility that BF and/or BF-shaped gut microbiota restricts the colonization of mucin degrading bacteria, particularly *A. muciniphila,* and *P. distasonis*, by suppressing mucin production in the distal gut. While BF itself has the ability to bind to and degrade mucin(61), the requirement of mucin for its colonization and persistence may not be as critical as that of other mucin degraders, particularly *A. muciniphila.* Of note, while mucin production is profoundly low in GF animals as compared to their conventional counterparts(66; 67), BF monocolonization alone did not alter *Muc2* expression significantly (not shown). Hence, whether BF or BF-shaped gut microbiota is responsible for inhibiting the mucin production pathway needs to be investigated in the future.

Our results appear to show that the ability of BF to restrict gut colonization by other gut microbes is more pronounced on *A. muciniphila*. In this regard, it has been shown that *A. muciniphila* colonization requires mucin for its O-glycan specific binding(59; 68; 69). With respect to the impact of distal gut competition by microbial communities, pre-clinical and clinical studies have shown that *A. muciniphila*, *P. distasonis,* and some members of the genus *Prevotella* are associated with protection from T1D(6; 17; 18; 70; 71), and they can synergistically protect from gut inflammation(72), suggesting that a persistent inhibitory effect on these communities by BF could result in accelerated autoimmune progression on T1D.

Importantly, it has been shown that variation in microbiome and LPS immunogenicity can impact T1D in humans(6). Immune response to *E. coli* LPS but not that of *Bacteroides* spp. Like *B. dorei* and *B. fragilis* is associated with early immune education and protection from T1D in young children(6; 22). One of these reports(6) also showed that, in addition to *E. coli*, diminished exposure to *A. muciniphila* is associated with higher T1D incidence in Finnish and Estonian children compared to Russian children. This study also showed that compared to *E.coli* and *A. muciniphila* LPS, polysaccharide preparations from many of the *Bacteroides* genera have a poor ability to induce cytokine production(6; 13). In agreement with these reports, not only do TLR4 knockout (KO) NOD mice exhibit a rapid onset of hyperglycemia (**5**), but also preparations of lipopolysaccharide (LPS) derived from Gram-negative bacteria, such as *Escherichia coli*, confer protection against T1D in NOD mice(73). However, our previous report(42) and current observations suggest an active role for TLR2, in addition to the influence of gut microbiota composition in BF-colonized mice and the potential differences in the LPS structure(6), in modulating the T1D disease outcomes. Our studies show that distal colon, spleen, and PnLN cells from BF-colonized mice of gnotobiotic and conventional facilities have relatively higher pro-inflammatory cytokine production and diminished expression of immune regulatory factors such as IL10, Foxp3, and IDO-1 as compared to controls.

Overall, higher abundance of Bacteroidetes phyla members including *Bacteroides* spp. has been detected not only in the gut of rodent models of T1D but also in T1D patients and at-risk children who have progressed towards developing disease symptoms(6; 12; 16-22), prompting the suggestion that some *Bacteroides* members could exert pro-autoimmune effects under T1D susceptibility. Our observations from this study, which examined the impact of juvenile age gut colonization by a commensal belonging to genus *Bacteroides* on T1D in NOD mice, show that gut commensals such as BF can contribute to initiation and/or perpetuation of autoimmunity in at-risk subjects. It is possible that early life gut colonization by BF contributes to two modes of perpetuation of inflammatory response in at risk backgrounds: 1) presence of BF in the gut at early ages can lead to TLR2-dependent inflammatory response in the systemic compartment due to underdeveloped epithelial barrier and higher gut permeability and 2) BF influences gut microbial community structure and function by actively suppressing colonization by commensal communities with the ability to promote immune regulation, gut integrity, and protection from T1D. Overall, this study sheds light on the potential mechanisms by which BF-like human gut commensals cause dysbiosis and impact disease outcomes in T1D prone subjects.

### Author contribution

R.G. researched data and edited the manuscript, H.T. researched and analyzed data, B.M.J researched data, R.M. researched data, M.E.M researched data, L.C. provided technical support, C.W. researched data and edited the manuscript, and C.V. designed experiments, researched, and analyzed data, and wrote the manuscript.

### Footnote

This work was supported by unrestricted research funds from MUSC and National Institutes of Health (NIH) grant R21AI133798 and R01DK136094. Dr. Vasu is the guarantor of this work and, as such, has full access to all the data in the study and takes responsibility for the integrity of the data and accuracy of the data analysis. The authors are thankful to Cell and Molecular Imaging, Pathology, immune monitoring and discovery, and flow cytometry cores of MUSC for the histology service, microscopy, FACS and multiplex assay instrumentation support.

## Abbreviations

TID: Type 1 Diabetes
NOD: non-obese diabetic
TLR: Toll-like receptor
PSA: polysaccharide A
BF: *Bacteroides fragilis*
PnLN: pancreatic lymph node
GF: germ free
NOD mice: non-obese diabetic mice.

## References

1. Charbonneau MR, Blanton LV, DiGiulio DB, Relman DA, Lebrilla CB, Mills DA, Gordon JI: A microbial perspective of human developmental biology. Nature 2016;535:48–55

2. Chervonsky AV: Microbiota and autoimmunity. Cold Spring Harbor perspectives in biology 2013;5:a007294

3. Arpaia N, Campbell C, Fan X, Dikiy S, van der Veeken J, deRoos P, Liu H, Cross JR, Pfeffer K, Coffer PJ, Rudensky AY: Metabolites produced by commensal bacteria promote peripheral regulatory T-cell generation. Nature 2013;504:451–455

4. Chiba T, Seno H: Indigenous clostridium species regulate systemic immune responses by induction of colonic regulatory T cells. Gastroenterology 2011;141:1114–1116

5. Wen L, Ley RE, Volchkov PY, Stranges PB, Avanesyan L, Stonebraker AC, Hu C, Wong FS, Szot GL, Bluestone JA, Gordon JI, Chervonsky AV: Innate immunity and intestinal microbiota in the development of Type 1 diabetes. Nature 2008;455:1109–1113

6. Vatanen T, Kostic AD, d’Hennezel E, Siljander H, Franzosa EA, Yassour M, Kolde R, Vlamakis H, Arthur TD, Hamalainen AM, Peet A, Tillmann V, Uibo R, Mokurov S, Dorshakova N, Ilonen J, Virtanen SM, Szabo SJ, Porter JA, Lahdesmaki H, Huttenhower C, Gevers D, Cullen TW, Knip M, Group DS, Xavier RJ: Variation in Microbiome LPS Immunogenicity Contributes to Autoimmunity in Humans. Cell 2016;165:842–853

7. Burrows MP, Volchkov P, Kobayashi KS, Chervonsky AV: Microbiota regulates type 1 diabetes through Toll-like receptors. Proceedings of the National Academy of Sciences of the United States of America 2015;112:9973–9977

8. The i MC: Household paired design reduces variance and increases power in multi-city gut microbiome study in multiple sclerosis. Mult Scler 2020:1352458520924594

9. i MCEasbue, i MC: Gut microbiome of multiple sclerosis patients and paired household healthy controls reveal associations with disease risk and course. Cell 2022;185:3467-3486 e3416

10. Azzouz DF, Chen Z, Izmirly PM, Chen LA, Li Z, Zhang C, Mieles D, Trujillo K, Heguy A, Pironti A, Putzel GG, Schwudke D, Fenyo D, Buyon JP, Alekseyenko AV, Gisch N, Silverman GJ: Longitudinal gut microbiome analyses and blooms of pathogenic strains during lupus disease flares. Ann Rheum Dis 2023;82:1315–1327

11. Vatanen T, Franzosa EA, Schwager R, Tripathi S, Arthur TD, Vehik K, Lernmark A, Hagopian WA, Rewers MJ, She JX, Toppari J, Ziegler AG, Akolkar B, Krischer JP, Stewart CJ, Ajami NJ, Petrosino JF, Gevers D, Lahdesmaki H, Vlamakis H, Huttenhower C, Xavier RJ: The human gut microbiome in early-onset type 1 diabetes from the TEDDY study. Nature 2018;562:589–594

12. Kostic AD, Gevers D, Siljander H, Vatanen T, Hyotylainen T, Hamalainen AM, Peet A, Tillmann V, Poho P, Mattila I, Lahdesmaki H, Franzosa EA, Vaarala O, de Goffau M, Harmsen H, Ilonen J, Virtanen SM, Clish CB, Oresic M, Huttenhower C, Knip M, Group DS, Xavier RJ: The dynamics of the human infant gut microbiome in development and in progression toward type 1 diabetes. Cell host & microbe 2015;17:260–273

13. Vatanen T, Kostic AD, d’Hennezel E, Siljander H, Franzosa EA, Yassour M, Kolde R, Vlamakis H, Arthur TD, Hamalainen AM, Peet A, Tillmann V, Uibo R, Mokurov S, Dorshakova N, Ilonen J, Virtanen SM, Szabo SJ, Porter JA, Lahdesmaki H, Huttenhower C, Gevers D, Cullen TW, Knip M, Group DS, Xavier RJ: Variation in Microbiome LPS Immunogenicity Contributes to Autoimmunity in Humans. Cell 2016;165:1551

14. Cho I, Yamanishi S, Cox L, Methe BA, Zavadil J, Li K, Gao Z, Mahana D, Raju K, Teitler I, Li H, Alekseyenko AV, Blaser MJ: Antibiotics in early life alter the murine colonic microbiome and adiposity. Nature 2012;488:621–626

15. Cox LM, Yamanishi S, Sohn J, Alekseyenko AV, Leung JM, Cho I, Kim SG, Li H, Gao Z, Mahana D, Zarate Rodriguez JG, Rogers AB, Robine N, Loke P, Blaser MJ: Altering the intestinal microbiota during a critical developmental window has lasting metabolic consequences. Cell 2014;158:705–721

16. Cinek O, Kramna L, Lin J, Oikarinen S, Kolarova K, Ilonen J, Simell O, Veijola R, Autio R, Hyoty H: Imbalance of bacteriome profiles within the Finnish Diabetes Prediction and Prevention study: Parallel use of 16S profiling and virome sequencing in stool samples from children with islet autoimmunity and matched controls. Pediatric diabetes 2016;

17. Alkanani AK, Hara N, Gottlieb PA, Ir D, Robertson CE, Wagner BD, Frank DN, Zipris D: Alterations in Intestinal Microbiota Correlate With Susceptibility to Type 1 Diabetes. Diabetes 2015;64:3510–3520

18. Mejia-Leon ME, Petrosino JF, Ajami NJ, Dominguez-Bello MG, de la Barca AM: Fecal microbiota imbalance in Mexican children with type 1 diabetes. Scientific reports 2014;4:3814

19. de Goffau MC, Luopajarvi K, Knip M, Ilonen J, Ruohtula T, Harkonen T, Orivuori L, Hakala S, Welling GW, Harmsen HJ, Vaarala O: Fecal microbiota composition differs between children with beta-cell autoimmunity and those without. Diabetes 2013;62:1238–1244

20. Sofi MH, Gudi R, Karumuthil-Melethil S, Perez N, Johnson BM, Vasu C: pH of drinking water influences the composition of gut microbiome and type 1 diabetes incidence. Diabetes 2014;63:632–644

21. de Goffau MC, Fuentes S, van den Bogert B, Honkanen H, de Vos WM, Welling GW, Hyoty H, Harmsen HJ: Aberrant gut microbiota composition at the onset of type 1 diabetes in young children. Diabetologia 2014;57:1569–1577

22. Davis-Richardson AG, Ardissone AN, Dias R, Simell V, Leonard MT, Kemppainen KM, Drew JC, Schatz D, Atkinson MA, Kolaczkowski B, Ilonen J, Knip M, Toppari J, Nurminen N, Hyoty H, Veijola R, Simell T, Mykkanen J, Simell O, Triplett EW: Bacteroides dorei dominates gut microbiome prior to autoimmunity in Finnish children at high risk for type 1 diabetes. Frontiers in microbiology 2014;5:678

23. Ley RE, Hamady M, Lozupone C, Turnbaugh PJ, Ramey RR, Bircher JS, Schlegel ML, Tucker TA, Schrenzel MD, Knight R, Gordon JI: Evolution of mammals and their gut microbes. Science 2008;320:1647–1651

24. Moore WE, Holdeman LV: Human fecal flora: the normal flora of 20 Japanese-Hawaiians. Appl Microbiol 1974;27:961–979

25. Nichols RL, Schumer W, Nythus LM, Bartlett JG, Gorbach SL: Anaerobic Infections. Am Fam Physician 1976;14:100–110

26. Bartlett JG, Onderdonk AB, Louie T, Kasper DL, Gorbach SL: A review. Lessons from an animal model of intra-abdominal sepsis. Arch Surg 1978;113:853–857

27. Round JL, Lee SM, Li J, Tran G, Jabri B, Chatila TA, Mazmanian SK: The Toll-like receptor 2 pathway establishes colonization by a commensal of the human microbiota. Science 2011;332:974–977

28. Onderdonk AB, Kasper DL, Cisneros RL, Bartlett JG: The capsular polysaccharide of Bacteroides fragilis as a virulence factor: comparison of the pathogenic potential of encapsulated and unencapsulated strains. The Journal of infectious diseases 1977;136:82–89

29. Liu CH, Lee SM, Vanlare JM, Kasper DL, Mazmanian SK: Regulation of surface architecture by symbiotic bacteria mediates host colonization. Proceedings of the National Academy of Sciences of the United States of America 2008;105:3951–3956

30. Kayama H, Takeda K: Polysaccharide A of Bacteroides fragilis: actions on dendritic cells and T cells. Molecular cell 2014;54:206–207

31. Dasgupta S, Erturk-Hasdemir D, Ochoa-Reparaz J, Reinecker HC, Kasper DL: Plasmacytoid dendritic cells mediate anti-inflammatory responses to a gut commensal molecule via both innate and adaptive mechanisms. Cell host & microbe 2014;15:413–423

32. Wang Q, McLoughlin RM, Cobb BA, Charrel-Dennis M, Zaleski KJ, Golenbock D, Tzianabos AO, Kasper DL: A bacterial carbohydrate links innate and adaptive responses through Toll-like receptor 2. The Journal of experimental medicine 2006;203:2853–2863

33. Chiu CC, Ching YH, Wang YC, Liu JY, Li YP, Huang YT, Chuang HL: Monocolonization of germ-free mice with Bacteroides fragilis protects against dextran sulfate sodium-induced acute colitis. Biomed Res Int 2014;2014:675786

34. Mazmanian SK, Liu CH, Tzianabos AO, Kasper DL: An immunomodulatory molecule of symbiotic bacteria directs maturation of the host immune system. Cell 2005;122:107–118

35. Cohen-Poradosu R, McLoughlin RM, Lee JC, Kasper DL: Bacteroides fragilis-stimulated interleukin-10 contains expanding disease. The Journal of infectious diseases 2011;204:363–371

36. Ochoa-Reparaz J, Mielcarz DW, Ditrio LE, Burroughs AR, Begum-Haque S, Dasgupta S, Kasper DL, Kasper LH: Central nervous system demyelinating disease protection by the human commensal Bacteroides fragilis depends on polysaccharide A expression. Journal of immunology 2010;185:4101–4108

37. Cameron G, Nguyen T, Ciula M, Williams SJ, Godfrey DI: Glycolipids from the gut symbiont Bacteroides fragilis are agonists for natural killer T cells and induce their regulatory differentiation. Chem Sci 2023;14:7887–7896

38. Ramakrishna C, Kujawski M, Chu H, Li L, Mazmanian SK, Cantin EM: Bacteroides fragilis polysaccharide A induces IL-10 secreting B and T cells that prevent viral encephalitis. Nature communications 2019;10:2153

39. Erturk-Hasdemir D, Oh SF, Okan NA, Stefanetti G, Gazzaniga FS, Seeberger PH, Plevy SE, Kasper DL: Symbionts exploit complex signaling to educate the immune system. Proceedings of the National Academy of Sciences of the United States of America 2019;116:26157–26166

40. Gautier T, Oliviero N, Ferron S, Le Pogam P, David-Le Gall S, Sauvager A, Leroyer P, Cannie I, Dion S, Sweidan A, Loreal O, Tomasi S, Bousarghin L: Bacteroides fragilis derived metabolites, identified by molecular networking, decrease Salmonella virulence in mice model. Frontiers in microbiology 2022;13:1023315

41. Hecht AL, Casterline BW, Earley ZM, Goo YA, Goodlett DR, Bubeck Wardenburg J: Strain competition restricts colonization of an enteric pathogen and prevents colitis. EMBO Rep 2016;17:1281–1291

42. Sofi MH, Johnson BM, Gudi RR, Jolly A, Gaudreau MC, Vasu C: Polysaccharide A-Dependent Opposing Effects of Mucosal and Systemic Exposures to Human Gut Commensal Bacteroides fragilis in Type 1 Diabetes. Diabetes 2019;68:1975–1989

43. Gudi R, Perez N, Johnson BM, Sofi MH, Brown R, Quan S, Karumuthil-Melethil S, Vasu C: Complex dietary polysaccharide modulates gut immune function and microbiota, and promotes protection from autoimmune diabetes. Immunology 2019;157:70–85

44. Perez N, Karumuthil-Melethil S, Li R, Prabhakar BS, Holterman MJ, Vasu C: Preferential costimulation by CD80 results in IL-10-dependent TGF-beta1(+) -adaptive regulatory T cell generation. Journal of immunology 2008;180:6566–6576

45. Karumuthil-Melethil S, Perez N, Li R, Vasu C: Induction of innate immune response through TLR2 and dectin 1 prevents type 1 diabetes. Journal of immunology 2008;181:8323–8334

46. Johnson BM, Gaudreau MC, Gudi R, Brown R, Gilkeson G, Vasu C: Gut microbiota differently contributes to intestinal immune phenotype and systemic autoimmune progression in female and male lupus-prone mice. Journal of autoimmunity 2020;108:102420

47. Taylor HB, Vasu C: Impact of Prebiotic beta-glucan Treatment at Juvenile Age on the Gut Microbiota Composition and the Eventual Type 1 Diabetes Onset in Non-obese Diabetic Mice. Front Nutr 2021;8:769341

48. Taylor HB, Gudi R, Brown R, Vasu C: Dynamics of Structural and Functional Changes in Gut Microbiota during Treatment with a Microalgal beta-Glucan, Paramylon and the Impact on Gut Inflammation. Nutrients 2020;12

49. Johnson BM, Gaudreau MC, Al-Gadban MM, Gudi R, Vasu C: Impact of dietary deviation on disease progression and gut microbiome composition in lupus-prone SNF1 mice. Clinical and experimental immunology 2015;181:323–337

50. Caporaso JG, Kuczynski J, Stombaugh J, Bittinger K, Bushman FD, Costello EK, Fierer N, Pena AG, Goodrich JK, Gordon JI, Huttley GA, Kelley ST, Knights D, Koenig JE, Ley RE, Lozupone CA, McDonald D, Muegge BD, Pirrung M, Reeder J, Sevinsky JR, Turnbaugh PJ, Walters WA, Widmann J, Yatsunenko T, Zaneveld J, Knight R: QIIME allows analysis of high-throughput community sequencing data. Nature methods 2010;7:335–336

51. Wixon J, Kell D: The Kyoto encyclopedia of genes and genomes--KEGG. Yeast 2000;17:48–55

52. Parks DH, Beiko RG: Identifying biologically relevant differences between metagenomic communities. Bioinformatics 2010;26:715–721

53. Hevia A, Milani C, Lopez P, Cuervo A, Arboleya S, Duranti S, Turroni F, Gonzalez S, Suarez A, Gueimonde M, Ventura M, Sanchez B, Margolles A: Intestinal dysbiosis associated with systemic lupus erythematosus. mBio 2014;5:e01548–01514

54. Zhang H, Liao X, Sparks JB, Luo XM: Dynamics of gut microbiota in autoimmune lupus. Applied and environmental microbiology 2014;80:7551–7560

55. Lu Y, Zhou G, Ewald J, Pang Z, Shiri T, Xia J: MicrobiomeAnalyst 2.0: comprehensive statistical, functional and integrative analysis of microbiome data. Nucleic Acids Res 2023;51:W310–W318

56. Ge SX, Son EW, Yao R: iDEP: an integrated web application for differential expression and pathway analysis of RNA-Seq data. BMC Bioinformatics 2018;19:534

57. Horn J, Bender BS, Bartlett JG: Role of anaerobic bacteria in perimandibular space infections. Ann Otol Rhinol Laryngol Suppl 1991;154:34–39

58. Polk BF, Kasper DL: Bacteroides fragilis subspecies in clinical isolates. Annals of internal medicine 1977;86:569–571

59. Wright DP, Rosendale DI, Robertson AM: Prevotella enzymes involved in mucin oligosaccharide degradation and evidence for a small operon of genes expressed during growth on mucin. FEMS Microbiol Lett 2000;190:73–79

60. Schwabkey ZI, Wiesnoski DH, Chang CC, Tsai WB, Pham D, Ahmed SS, Hayase T, Ortega Turrubiates MR, El-Himri RK, Sanchez CA, Hayase E, Frenk Oquendo AC, Miyama T, Halsey TM, Heckel BE, Brown AN, Jin Y, Raybaud M, Prasad R, Flores I, McDaniel L, Chapa V, Lorenzi PL, Warmoes MO, Tan L, Swennes AG, Fowler S, Conner M, McHugh K, Graf T, Jensen VB, Peterson CB, Do KA, Zhang L, Shi Y, Wang Y, Galloway-Pena JR, Okhuysen PC, Daniel-MacDougall CR, Shono Y, Burgos da Silva M, Peled JU, van den Brink MRM, Ajami N, Wargo JA, Reddy P, Valdivia RH, Davey L, Rondon G, Srour SA, Mehta RS, Alousi AM, Shpall EJ, Champlin RE, Shelburne SA, Molldrem JJ, Jamal MA, Karmouch JL, Jenq RR: Diet-derived metabolites and mucus link the gut microbiome to fever after cytotoxic cancer treatment. Sci Transl Med 2022;14:eabo3445

61. Huang JY, Lee SM, Mazmanian SK: The human commensal Bacteroides fragilis binds intestinal mucin. Anaerobe 2011;17:137–141

62. Macfarlane GT, Gibson GR: Formation of glycoprotein degrading enzymes by Bacteroides fragilis. FEMS Microbiol Lett 1991;61:289–293

63. Chatzidaki-Livanis M, Coyne MJ, Comstock LE: An antimicrobial protein of the gut symbiont Bacteroides fragilis with a MACPF domain of host immune proteins. Mol Microbiol 2014;94:1361–1374

64. Coyne MJ, Roelofs KG, Comstock LE: Type VI secretion systems of human gut Bacteroidales segregate into three genetic architectures, two of which are contained on mobile genetic elements. BMC Genomics 2016;17:58

65. Coyne MJ, Bechon N, Matano LM, McEneany VL, Chatzidaki-Livanis M, Comstock LE: A family of anti-Bacteroidales peptide toxins wide-spread in the human gut microbiota. Nature communications 2019;10:3460

66. Kandori H, Hirayama K, Takeda M, Doi K: Histochemical, lectin-histochemical and morphometrical characteristics of intestinal goblet cells of germfree and conventional mice. Exp Anim 1996;45:155–160

67. Enss ML, Grosse-Siestrup H, Schmidt-Wittig U, Gartner K: Changes in colonic mucins of germfree rats in response to the introduction of a "normal" rat microbial flora. Rat colonic mucin. J Exp Anim Sci 1992;35:110–119

68. Van Herreweghen F, Van den Abbeele P, De Mulder T, De Weirdt R, Geirnaert A, Hernandez-Sanabria E, Vilchez-Vargas R, Jauregui R, Pieper DH, Belzer C, De Vos WM, Van de Wiele T: In vitro colonisation of the distal colon by Akkermansia muciniphila is largely mucin and pH dependent. Benef Microbes 2017;8:81–96

69. Elzinga J, Narimatsu Y, de Haan N, Clausen H, de Vos WM, Tytgat HLP: Binding of Akkermansia muciniphila to mucin is O-glycan specific. Nature communications 2024;15:4582

70. Hansen CH, Krych L, Nielsen DS, Vogensen FK, Hansen LH, Sorensen SJ, Buschard K, Hansen AK: Early life treatment with vancomycin propagates Akkermansia muciniphila and reduces diabetes incidence in the NOD mouse. Diabetologia 2012;55:2285–2294

71. Hanninen A, Toivonen R, Poysti S, Belzer C, Plovier H, Ouwerkerk JP, Emani R, Cani PD, De Vos WM: Akkermansia muciniphila induces gut microbiota remodelling and controls islet autoimmunity in NOD mice. Gut 2018;67:1445–1453

72. Gaifem J, Mendes-Frias A, Wolter M, Steimle A, Garzon MJ, Ubeda C, Nobre C, Gonzalez A, Pinho SS, Cunha C, Carvalho A, Castro AG, Desai MS, Rodrigues F, Silvestre R: Akkermansia muciniphila and Parabacteroides distasonis synergistically protect from colitis by promoting ILC3 in the gut. mBio 2024;15:e0007824

73. Caramalho I, Rodrigues-Duarte L, Perez A, Zelenay S, Penha-Goncalves C, Demengeot J: Regulatory T cells contribute to diabetes protection in lipopolysaccharide-treated non-obese diabetic mice. Scandinavian journal of immunology 2011;74:585–595

