## Supplemental Figures 1-3 for "Gut colonization by *Bacteroides fragilis* at juvenile age alters microbiota composition and accelerates type 1 diabetes progression in non-obese diabetic mice"

**Fig. S1**

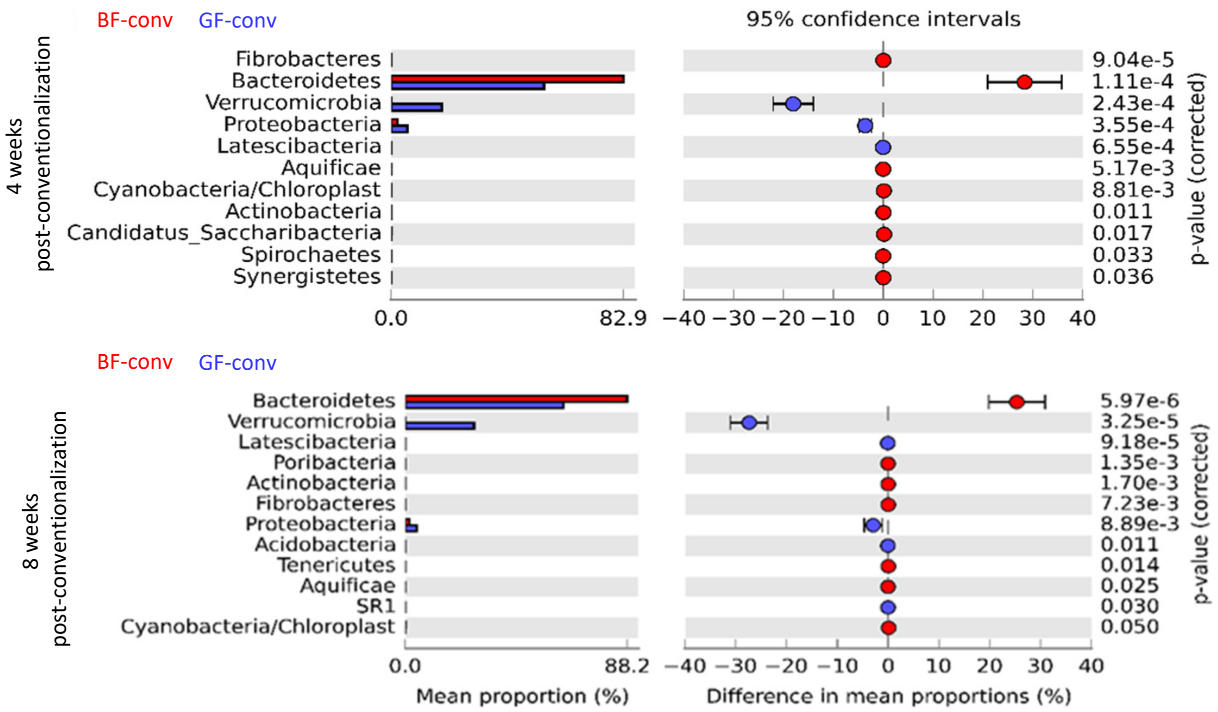


***Figure S1: Impact of BF on the acquisition and overall community profiles of gut microbiota.*** Fecal pellets were collected from conventionalized GF (GF-conv / Ex-GF) and conventionalized BF-monocolonized (BF-conv / Ex-BF) mice at 4- and 8-weeks post-conventionalization, and the DNA preparations were subjected to 16S rRNA gene (V3/V4 region) -targeted sequencing using the Illumina MiSeq platform and the sequence data was analyzed as described for Figs. 2 and 6. Mean relative abundances of sequences representing microbial communities at phylum level are shown. Statistical analysis: two-sided Welch's t-test and the *p*-values were FDR corrected using Benjamini and Hochberg approach.

**Fig. S2**

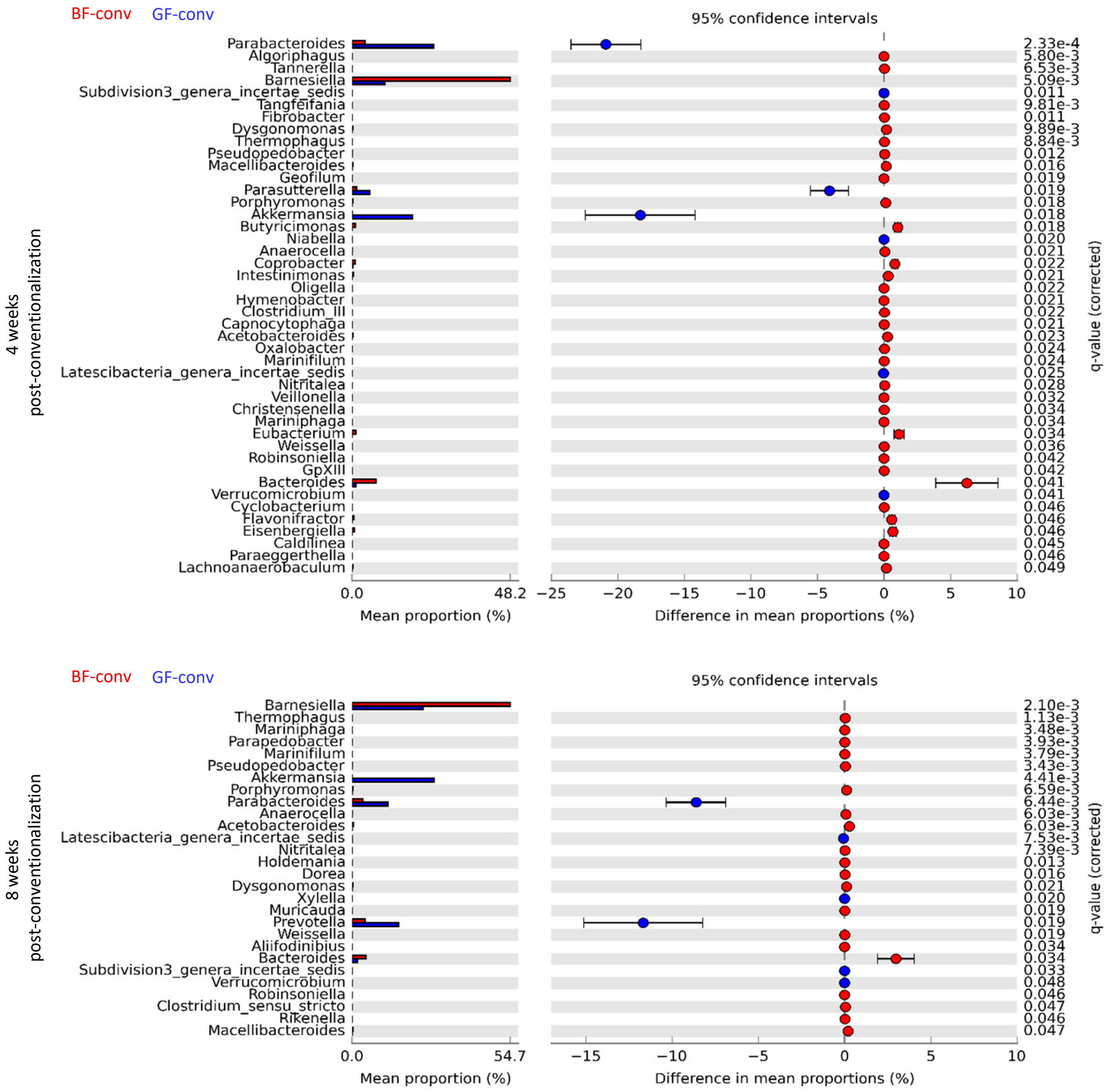


***Figure S1: Impact of BF on the acquisition and overall community profiles of gut microbiota.*** Fecal pellets were collected from conventionalized GF (GF-conv / Ex-GF) and conventionalized BF-monocolonized (BF-conv / Ex-BF) mice at 4- and 8-weeks post-conventionalization, and the DNA preparations were subjected to 16S rRNA gene (V3/V4 region) -targeted sequencing using the Illumina MiSeq platform and the sequence data was analyzed as described for Figs. 2 and 6. Mean relative abundances of sequences representing microbial communities at genus level are shown. Statistical analysis: two-sided Welch's t-test and the *p*-values were FDR corrected using Benjamini and Hochberg approach.

**Fig. S2**

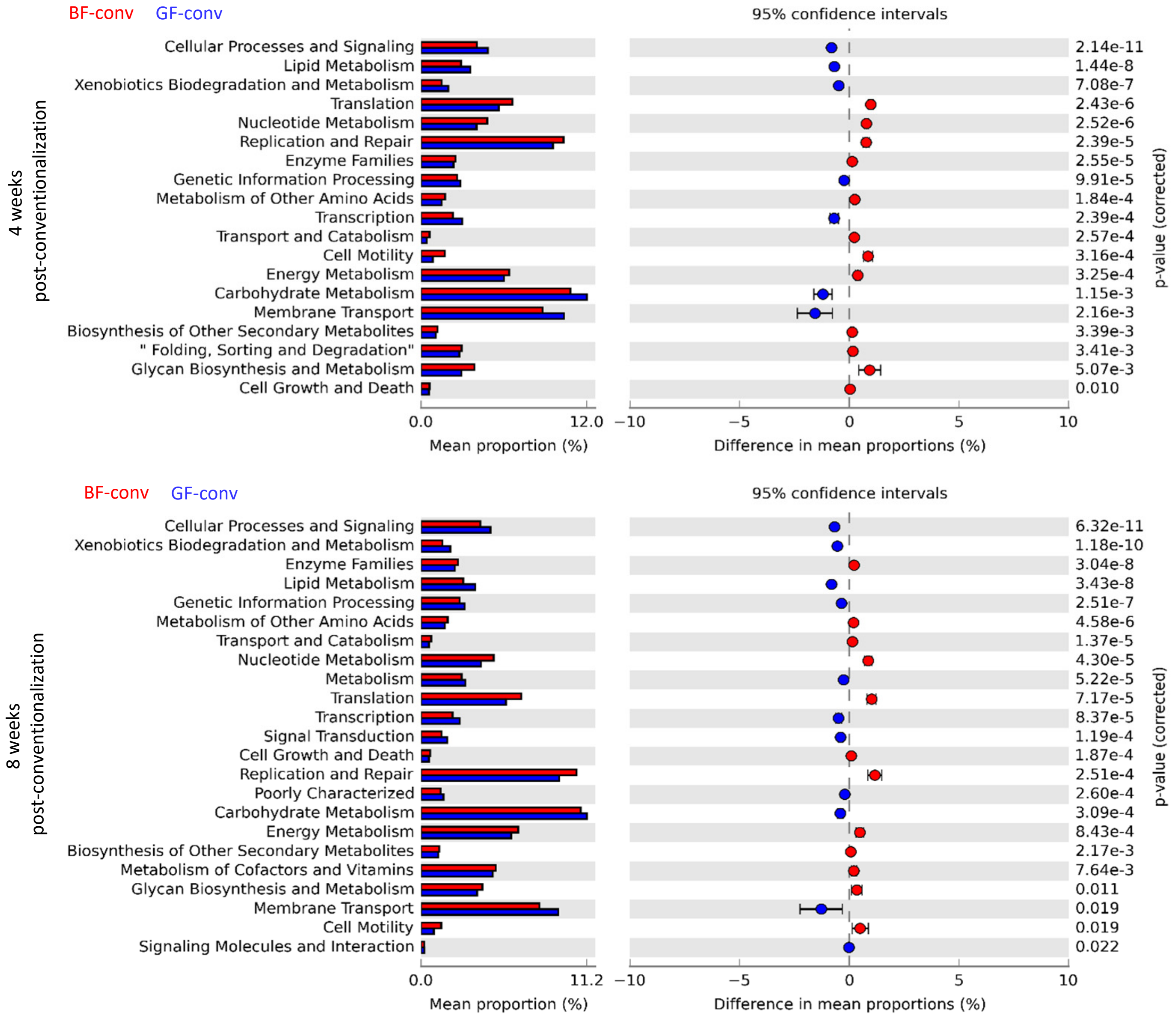


***Figure S2: Impact of BF on the overall predictive functional profiles of fecal microbiota.*** Fecal pellets were collected from conventionalized GF (GF-conv / Ex-GF) and conventionalized BF-monocolonized (BF-conv / Ex-BF) mice at 4- and 8-weeks post-conventionalization, and the DNA preparations were subjected to 16S rRNA gene (V3/V4 region) -targeted sequencing using the Illumina MiSeq platform as described for Figs. 2 and 6. Sequencing data was used for metagenomes prediction of KEGG orthologs employing PICRUSt application and the level 2 predictive emetabolic function data was visualized using STAMP. Statistical analysis: two-sided Welch's t-test and the *p*-values were FDR corrected using Benjamini and Hochberg approach.
